# In silico identification of novel chemical compounds with anti-TB potential for the inhibition of InhA and EthR from Mycobacterium tuberculosis

**DOI:** 10.1101/2020.12.04.411967

**Authors:** Sajal Kumar Halder, Fatiha Elma

## Abstract

Tuberculosis (TB) continuously pose a major public health concern around the globe, with a mounting death toll of approximately 1.4 million in 2019. The reduced bioavailability, increased toxicity and resistance of several first-line and second-line anti-TB drugs such as isoniazid, ethionamide have necessitated the search for new medications. In this research, we have identified several novel chemical compounds with anti-TB properties using various computational tools like molecular docking analysis, drug-likeness evaluation, ADMET profiling, P450 site of metabolism prediction and molecular dynamics simulation study. This study involves fifty drug-like compounds with antibacterial activity that inhibit InhA and EthR involved in the synthesis of one of the major lipid components, mycolic acid, which is crucial for the viability of *Mycobacterium tuberculosis.* Among these fifty compounds, 3-[3-(4-Fluorophenyl)-1,2,4-oxadiazol-5-yl]-N-(2-methylphenyl) piperidine-1-carboxamide (C22) and 5-(4-Ethyl-phenyl)-2-(1H-tetrazol-5-ylmethyl)-2H-tetrazole (C29) were found to pass the two-step molecular docking, P450 site of metabolism prediction and pharmacokinetics filtering analysis successfully. Their binding stability for target proteins have been evaluated through RMSD, RMSF, Radius of gyration analysis from 10 ns Molecular Dynamics Simulation (MDS) run. Our identified drugs could be a capable therapeutic for Tuberculosis drug discovery, having said that more in vitro and in vivo testing is required to justify their potential as novel drug and mode of action.

## 1. INTRODUCTION

Tuberculosis (TB) is a transmissible disease brought about by *Mycobacterium tuberculosis* (Mtb), which brings severe damage to lungs (pulmonary TB), along with other regions of the body, i.e. brain, spinal cord, lymph node, abdomen, bones/joints, intestinal and genitourinary system (extra pulmonary TB) [1, 2, 3]. Active TB symptoms depends on the severity of spread, generally some of clinical features are: persistent coughing more than 3 weeks, bloody sputum (hemoptysis), chest pain, fever, fatigue/weakness, weight loss, anorexia, breathlessness etc. The person with latent tuberculosis infection (LTBI) presents no symptoms and cannot transmit Mtb to others, but may develop active TB [4]. WHO recommended a standard 6-month administrative treatment for active, pulmonary, and pharmaco-sensitive TB with different combinations of four first-line medications: isoniazid, rifampin, ethambutol and pyrazinamide as bases for treatment [2, 7]. Although TB is a treatable infection, the massive emergence of resistance to antibiotics makes it a global threat. The classes of resistance that are becoming more and more prevalent includes (i) RR-TB, resistance to the first line drug rifampin only (ii) MDR-TB or multidrug resistance TB defines as the resistance towards at least two first line drugs, isoniazid and rifampin (iii) XDR-TB or extensively drug resistance TB defines as MDR-TB combined with resistance to a minimum of one drug from every of the two classes of second-line drugs (fluoroquinolones and injectables) and (iv) TDR-TB or totally drug resistance defines as the resistance to more drugs than strains categorized as XDR-TB [5,6].

It is undeniable that, the world needs new drugs to improve current treatment of TB to get around the escalation of drug-resistance. Therefore, our in-silico approach seeks to offset the burden of existing drug resistance scenarios of through exploration of new drugs performance. As a part of the experiment, a total fifty drug-like compounds were collected to assess their effectiveness in the treatment of TB and clinical management. Our experiment proceeded through evaluating binding affinity of fifty ligands utilizing two different docking tools DockThor and Autodock vina individually against the InhA (PDB ID: 3FNG) and EthR (PDB ID: 3G1M) receptor proteins. Later on, leading four compounds were analyzed for predicting probable metabolic sites of CYP. The study was further validated by assessing drug-likeness properties and ADMET prediction so as to examine their pharmacokinetics and pharmacodynamics properties. Then the best compounds with receptor complex were subjected to molecular dynamics simulations to test the thermodynamic properties.

### 1.1. InhA (Enoyl acyl carrier protein reductase), a pivotal enzyme used in Mycolic acid biosynthesis pathway

Mycobacteria are distinctive in having two Fatty acid synthase (FAS) systems which ends up synthesizing the mycobacterial mycolic acid (MA) precursors [8]: the “eukaryotic-type” multifunctional FAS-I for de novo synthesis of long-chain fatty acyl-CoA and their subsequent elongation by the “bacterial-type” FAS-II [9,10]. Malonyl-CoA is produced by the carboxylation of acetyl-CoA catalyzed by acetyl-CoA carboxylase (ACC) and integrated into the developing acylic chain during the repeated cycle of reactions of fatty acids synthase I and II (FAS I/FAS II) [11]. The mero-mycolic chain precursors are synthesized in four stages of iterative cycles with four FAS-II enzymes catalyzing acyl carrier protein (ACP) derivatives in each elongation cycle [8]. The transformation of malonyl-CoA to malonyl-ACP by malonyl-CoA:ACP transacylase (mtFabD) facilitates its integration into the FAS-II enzymatic system [12]. Both FAS systems are interconnected by a CoA-dependent linker enzyme β-ketoacyl-ACP synthase III (mtFabH) which catalyzes Claisen condensation of acyl-CoA precursors produced by FAS-I and malonyl-ACP to form β-ketoacyl-ACPs [13]. Next, β-ketoacyl-ACP reductase (MabA) performs NADP+H^+^-specific reduction of long chain β-ketoacyl-ACP resulting in β-hydroxyacyl-ACP intermediate which is dehydrated by the β-hydroxyacyl-ACP dehydratase complex (HadABC) to an enoyl-ACP [14,15]. The end step of the elongation cycle involves trans-2-enoyl-ACP reductase (InhA), which catalyzes the reduction of NADH+H^+^ in the enoyl chain and develop long-chain saturated acyl-ACP [16]. β-ketoacyl-ACP synthases (KasA and KasB) catalyzes new acyl extension cycles with a new malonyl-ACP unit, thereby extending the growing mero-mycolate chain by two carbon units to form a β-ketoacyl-ACP from acyl-ACP and malonyl-ACP [17,18]. Finally, last Claisen-type condensation step catalyzed by polyketide synthase (Pks13) that connects the mero-mycolic chain to a carboxylated α-chain produced by FAS-I that synthesizes trehalose mycolic β-ketoester and reduced by CmrA to trehalose monomycolate (TMM), a form of mature mycolic acid in Mtb [19,20,21]. The detailed biosynthesis pathway of mycolic acid is displayed in Figure 1.

**Figure 1:**
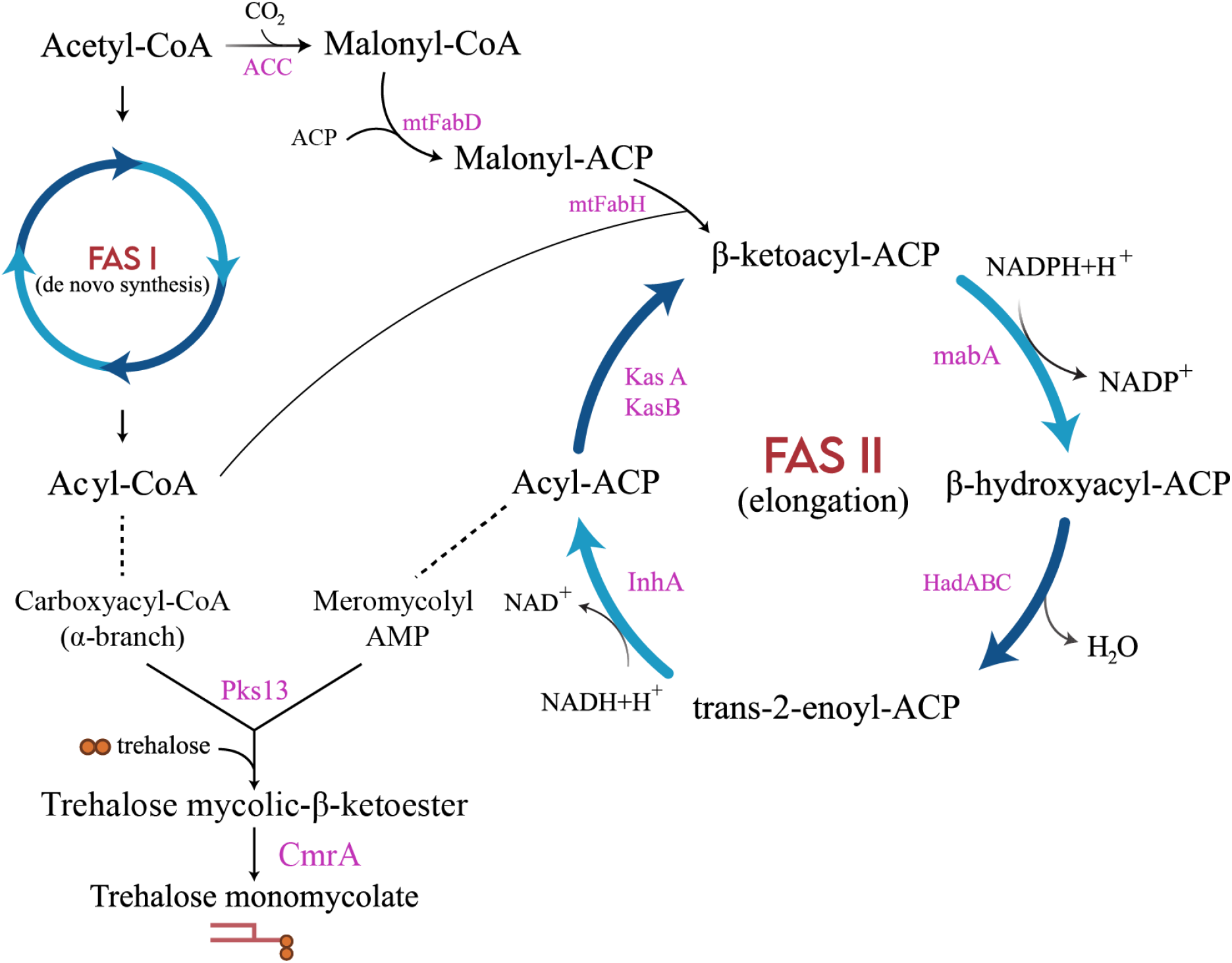
Biosynthesis of MA in Mtb. The acetyl-CoA carboxylase (ACC) produces malonyl-CoA, later on the synthesis of malonyl-ACP occurs by malonyl-CoA:AcpM transacylase (mtFabD). FAS II is triggered by the β-ketoacyl-ACP synthase (mtFabH), which condenses the acyl-CoA to malonyl-ACP to generate a β-ketoacyl-ACP. Following the condensation stage, the β-ketoacyl-ACP passes through a series of keto-reduction, dehydration, and enoyl-reduction catalyzed by the β-ketoacyl-ACP reductase (MabA), the β-hydroxyacyl-ACP dehydratase complex (HadABC) and the enoyl-ACP reductase (InhA), respectively. Further elongation cycles are initiated either by KasA or KasB β-ketoacyl ACP synthase. The last condensation carried out by polyketide synthase (Pks13) delivers trehalose mycolic β-ketoester and then reduced to trehalose monomycolate (TMM) [ADAPTED FROM 19].

### 1.2. Proposed anti-TB drugs acting against InhA and EthR

In our study, the promising targets of two of our selected ligands, 3-[3-(4-Fluorophenyl)-1,2,4-oxadiazol-5-yl]-N-(2-methylphenyl) piperidine-1-carboxamide and 5-(4-Ethyl-phenyl)-2-(1H-tetrazol-5-ylmethyl)-2H-tetrazole were InhA and EthR respectively. Both inhibitors are offered to work against InhA and EthR accordingly, interfering with the biosynthetic pathway of MA. InhA participates in the synthesis of MA, a major constituent of the mycobacterial cell wall [16] and EthR belongs to the tetR/CamR family of transcriptional repressor that negatively regulates the expression of EthA enzyme (Figure 2) [22]. INH and ETH both act as prodrug and represent metabolic activation dependency within Mtb cells by two mycobacterial enzymes, catalase peroxidase katG and FAD containing monooxygenase EthA respectively [23,24]. Both require the formation of covalent adducts with NAD co-factor on oxidative activation to act on the InhA enzyme which is involved in mycolic acid synthesis. (Figure 2) [25,26,27].

**Figure 2:**
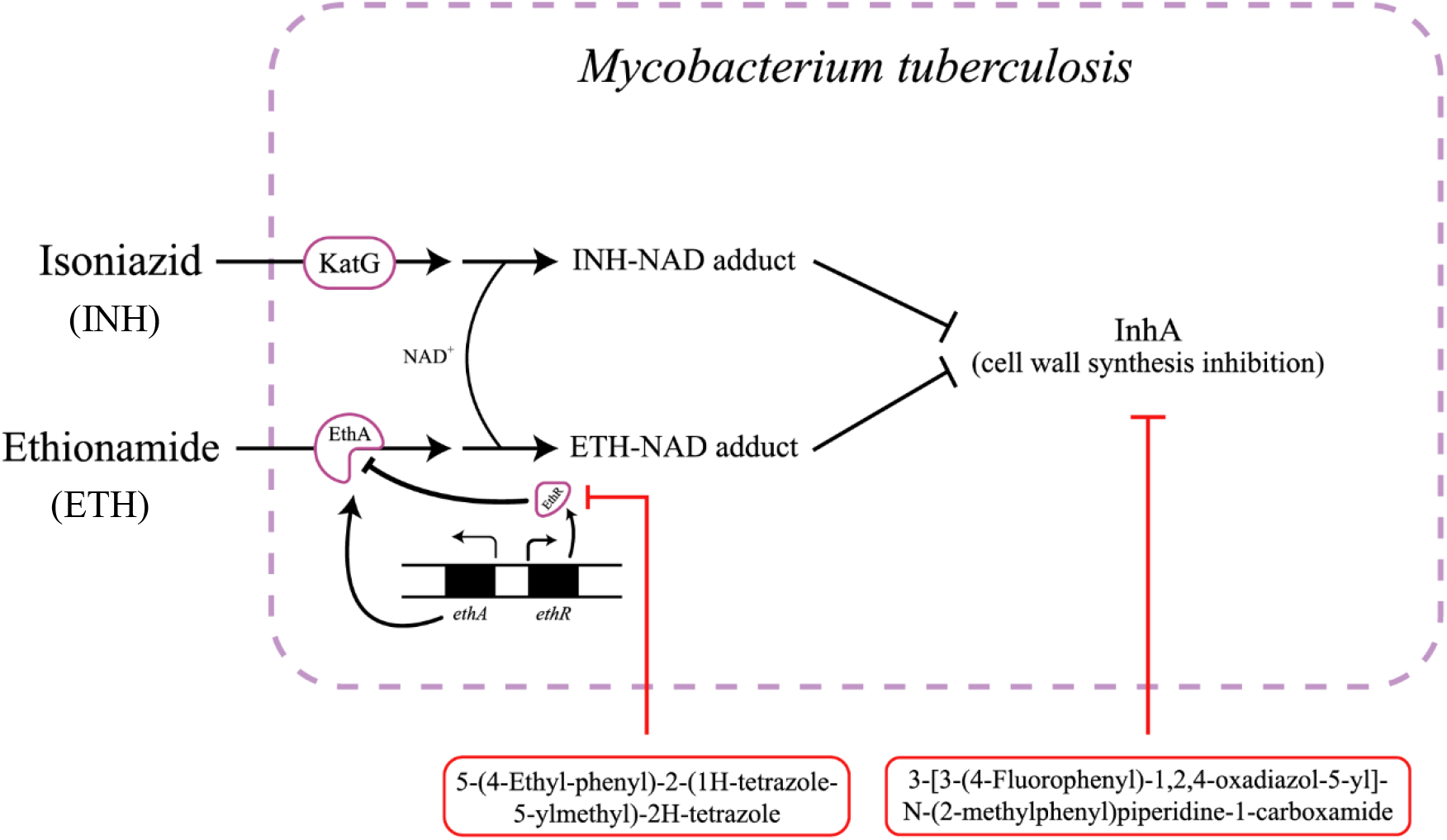
A schematic diagram showing mechanism of action of selected control drugs, Isoniazid (INH) and Ethionamide (ETH) along with showing proposed novel anti-TB drugs, 3-[3-(4-Fluorophenyl)-1,2,4-oxadiazol-5-yl]-N-(2-methylphenyl) piperidine-1-carboxamide and 5-(4-Ethyl-phenyl)-2-(1H-tetrazol-5-ylmethyl)-2H-tetrazole acting against InhA and EthR inside the Mtb. Both prodrugs, INH and ETH get metabolically activated by katG and EthA respectively, continuing towards formation of NAD adducts with their respective activated forms which eventually inhibit the enzyme InhA involved in mycolic acid biosynthesis. EthR, a negative transcriptional regulator which reduces the expression of EthA enzyme causing the blockage of the ETH activation pathway. Our selected two novel inhibitors, 3-[3-(4-Fluorophenyl)-1,2,4-oxadiazol-5-yl]-N-(2-methylphenyl) piperidine-1-carboxamide and 5-(4-Ethyl-phenyl)-2-(1H-tetrazol-5-ylmethyl)-2H-tetrazole are proposed to act against InhA and EthR respectively, leading to disrupting the MA biosynthesis pathway. [ADAPTED FROM: 26,28]

## 2. MATERIALS AND METHODS

### 2.1. Molecular docking

#### 2.1.1. Initial Molecular Docking in DockThor program

The preliminary docking of two receptor protein and fifty ligands were carried out by DockThor docking server (https://www.dockthor.lncc.br/v2/). The algorithm of the program is based on flexible-ligand and rigid-receptor grid-based system. This web-based tool spontaneously produces the topology files (i.e., atom types and partial charges) for the protein, ligand, and cofactors according to the MMFF94S force field [29]. The crystal structures of two receptors i.e., Crystal structure of InhA bound to triclosan derivative (PDB ID: 3FNG) and EthR from Mycobacterium tuberculosis in complex with compound BDM31381 (PDB ID: 3G1M), were taken from Protein Data Bank (https://www.rcsb.org/search). Fifty compounds were collected from both ACD: Antibacterial Chemotherapeutics Database (http://amdr.amu.ac.in/acd/index.jsp) and PubChem (https://pubchem.ncbi.nlm.nih.gov/) servers. Isoniazid (INH) and Ethionamide (ETH) were used as control drug against InhA and EthR proteins respectively. In the beginning, the PDB structures were modified using PyMOL tools (PyMOL) by clearing the water molecules from the structure [30] and then minimizing the structure employing Swiss-PdbViewer [31].

#### 2.1.2. Autodock-vina binding affinity prediction

Following the initial affinity prediction, desired ligands with lower affinity score than control drugs were chosen to evaluate bound conformation and binding energy. Autodock vina software (AutoDock 4.2) was used to evaluate bound conformation and binding energy of those selected ligands [32]. This tool uses Lamarckian Genetic Algorithm to evaluate binding energy of ligands with receptor proteins. Exhaustiveness defines how many times the system will repeat the computation on binding site of protein and this was kept 10 for the analysis. CASTp server [33] was used to identify ligand binding sites of proteins and prepare grid box.

### 2.2. Visualization and interaction analysis

The 2D and 3D visualization of non-bonded interactions of protein-ligand docked complexes were done by BIOVIA Discovery Studio 4.1 Visualizer [34]. This tool was utilized to get the total number of hydrogen bonds and interacting amino acids of proteins.

### 2.3. P450 Site of Metabolism (SOM) prediction

The most promising four compounds were chosen for further calculation. The probable sites of metabolism of these compounds were predicted using RS-Web Predictor 1.0 (http://reccr.chem.rpi.edu/Software/RSWebPredictor/) [35]. This prediction tool uses 3A4, 1A2, 2A6, 2B6, 2C8, 2C9, 2D6, 2E1, 3A4 CYP isoforms for cytochrome P450 site evaluation.

### 2.4. Drug-likeness Properties analysis and ADMET prediction

Determination of drug likeness property of drug-like compounds is one of the vital steps of drug discovery. These properties were predicted utilizing Lipinski’s rule of five [36], Ghose’s rule [37] Veber’s rule [38] Muegge’s rule [39], TPSA and No of rotatable bonds. The calculation was carried out using SwissADME online tool (http://www.swissadme.ch/index.php) [40].

Absorption, distribution, metabolism, excretion and toxicity, all these chemical properties were also crucial determinant of drug like compounds. ADMET properties of each of four compounds were predicted using another online tool, admetSAR (http://lmmd.ecust.edu.cn/admetsar2/) [41]. In each of the prediction study, canonical smiles of the compounds were used from PubChem database (https://pubchem.ncbi.nlm.nih.gov/).

### 2.5. Molecular Dynamics Simulation

The study of molecular dynamics simulation (MDS) is a thermodynamics-based operation which helps to study the dynamic perturbation found in protein-ligand complexes. In our experiment, to ensure the stability of protein-ligand complex, we subjected the best ligands screened from previous steps to the molecular dynamics simulation (MDS) study with their respective proteins. We simulated the docking complexes using the NAMD_2.14bNAMD_2.14b2_Win64-multicore-CUDA version [42] implying CHARMM 36 force field [43] and TIP3P water model. A multi-step time algorithm was used, with an integration time step of 2 femto second. Visual molecular dynamics (VMD) [44] was used to generate psf files of protein-ligand complexes, water box and for neutralizing the system with sodium (Na+) and a chloride (Cl-) ions. Ligand topology and parameter files were generated using CHARMM-GUI web service [45]. The simulation was run for 10 ns, where the system was minimized for 1000 steps. Langevin thermostat was used to maintain a constant temperature of 310k. Periodic boundary conditions was applied surrounding the system. Finally, the results were visualized and analyzed in VMD.

## 3. RESULTS

### 3.1. Molecular docking analysis

To filter out the best ligands from docking analysis, two separate docking tools were used with different algorithm. Initially used DockThor (server-based docking tool) results were further scanned through Autodock-vina software. Detailed information of all the collected drug-like compounds were provided in *Supplementary material.*

Total fifty ligands were primarily docked individually against the InhA (PDB ID: 3fng) and EthR (PDB ID: 3g1m) receptors protein via the DockThor server. Against both proteins, InhA and EthR, all the compounds showed better affinity than control 1 (Isoniazid) and control 2 (Ethionamide). The binding affinity score between InhA, EthR receptors and fifty ligands, as well as control drugs is depicted in Table 1.

**Table 1:**
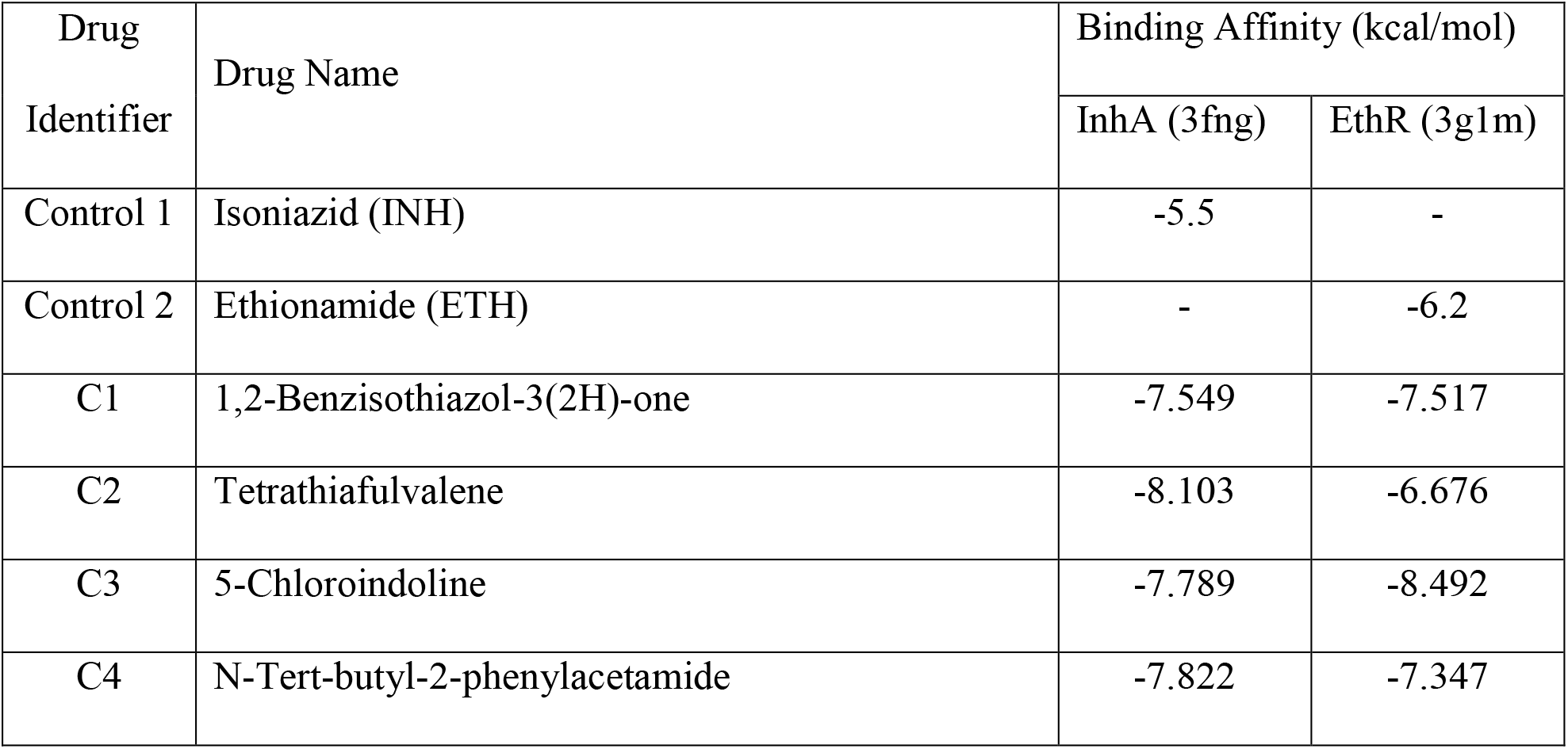
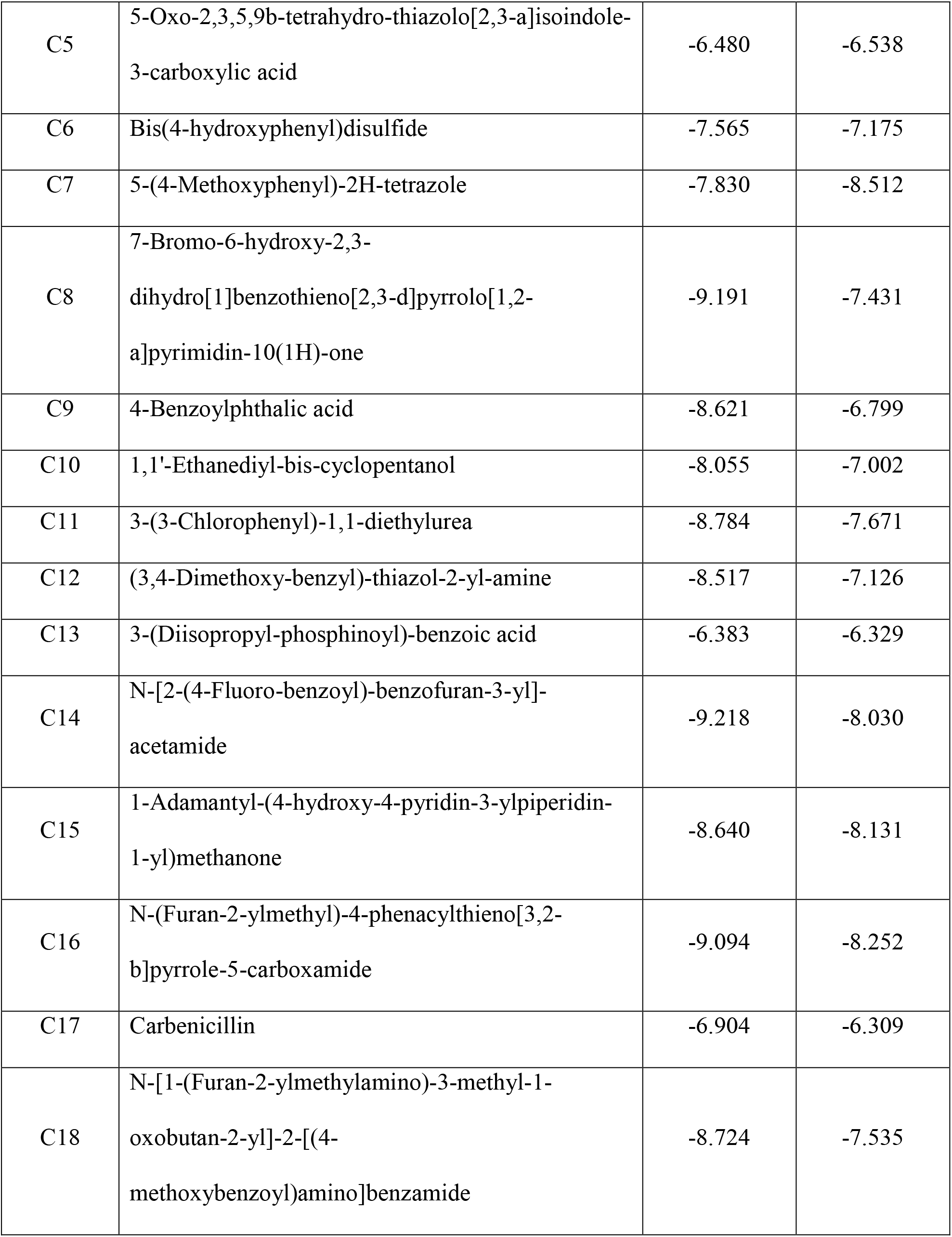
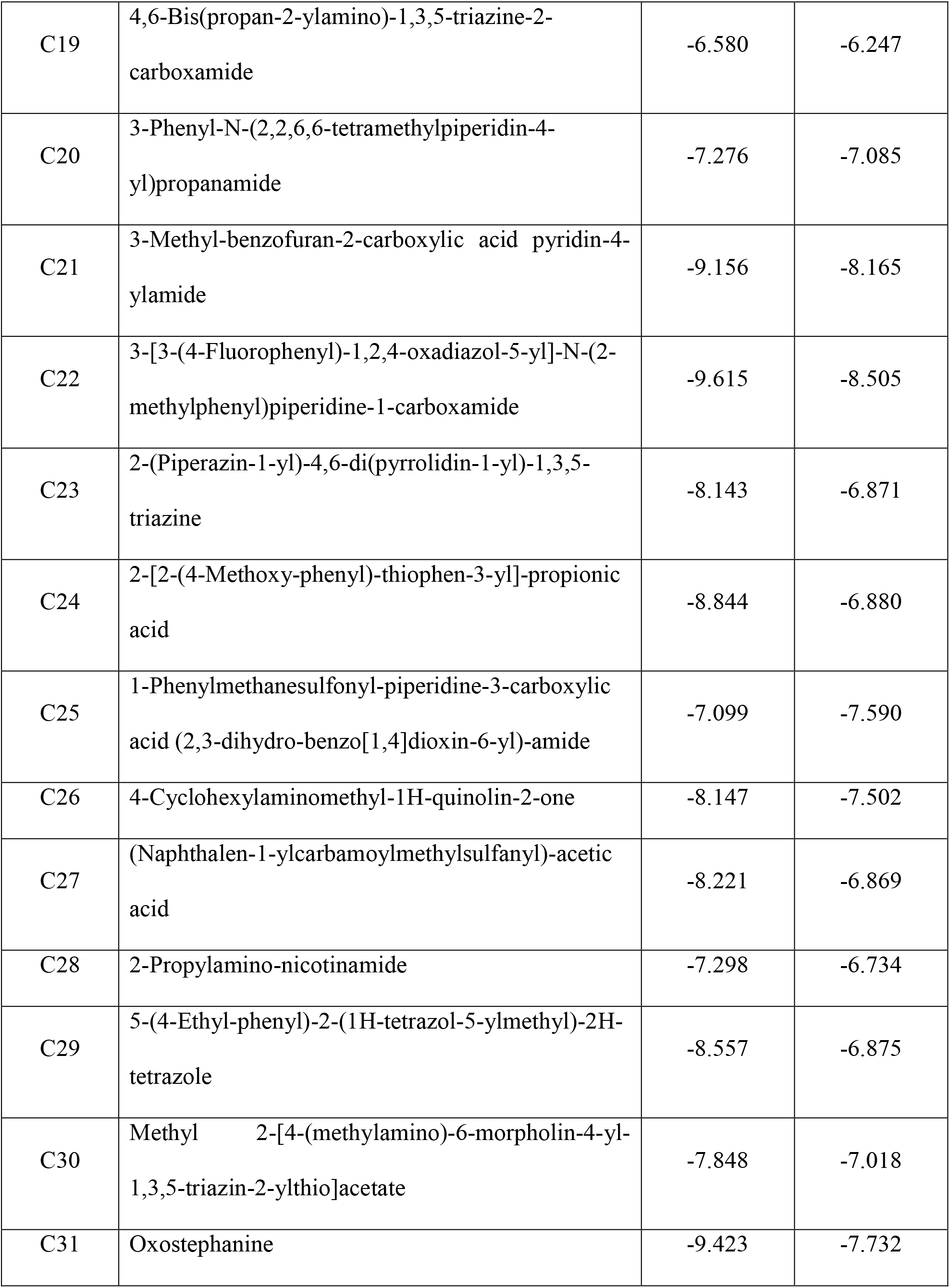
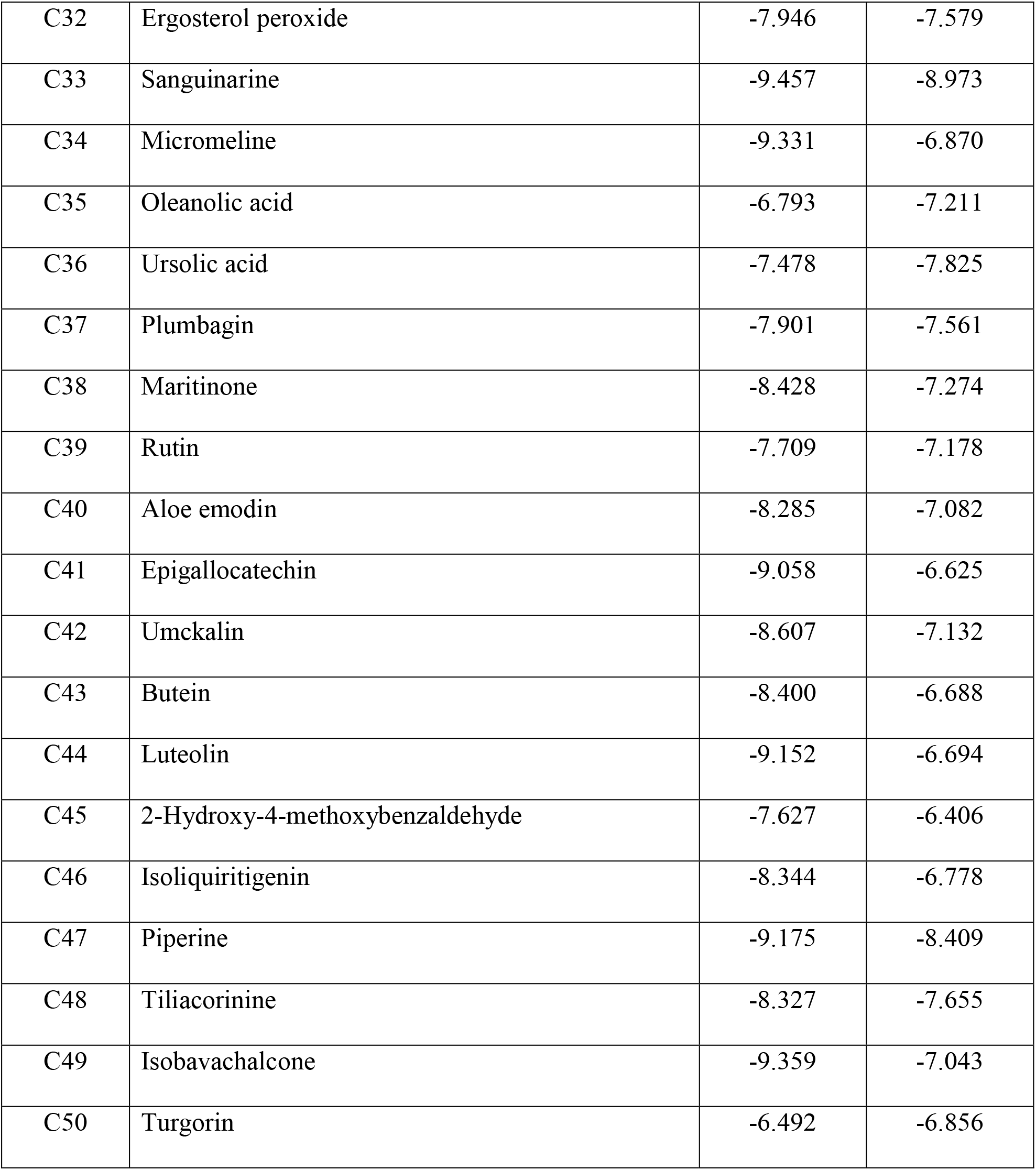
Binding affinity score between receptors (InhA, EthR) and fifty ligands, along with control drugs.

Ligands, filtered by DockThor server were considered for computing binding energy and bound conformation with the assistance of Autodock Vina software. Two ligands were preferred over all ligands considering their lowest binding score towards InhA protein. Likewise, two ligands with lowest binding score towards EthR protein were chosen for further analysis. Comparative Docking results of Autodock-vina tool are listed in Table 2.

**Table 2:**
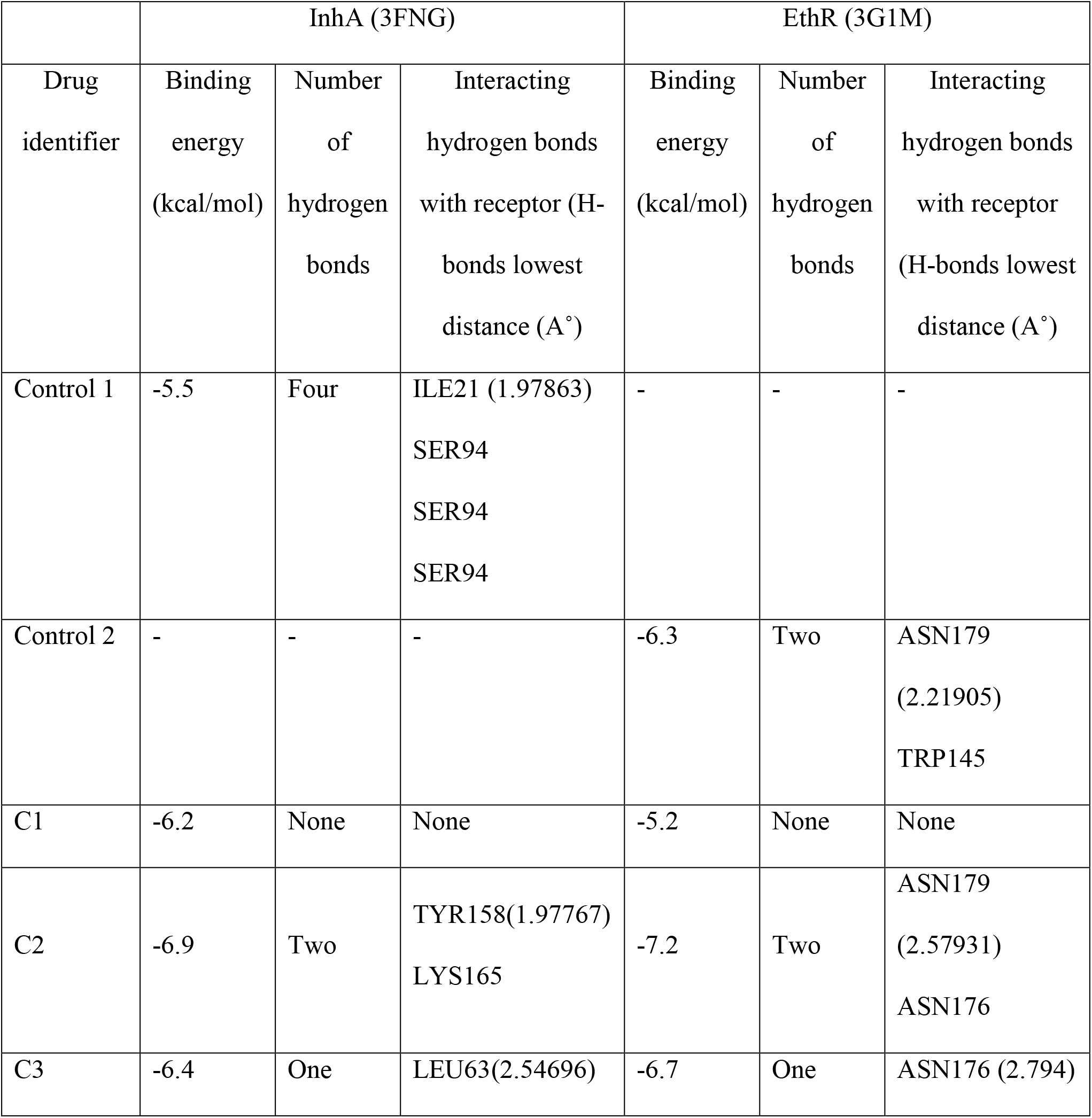
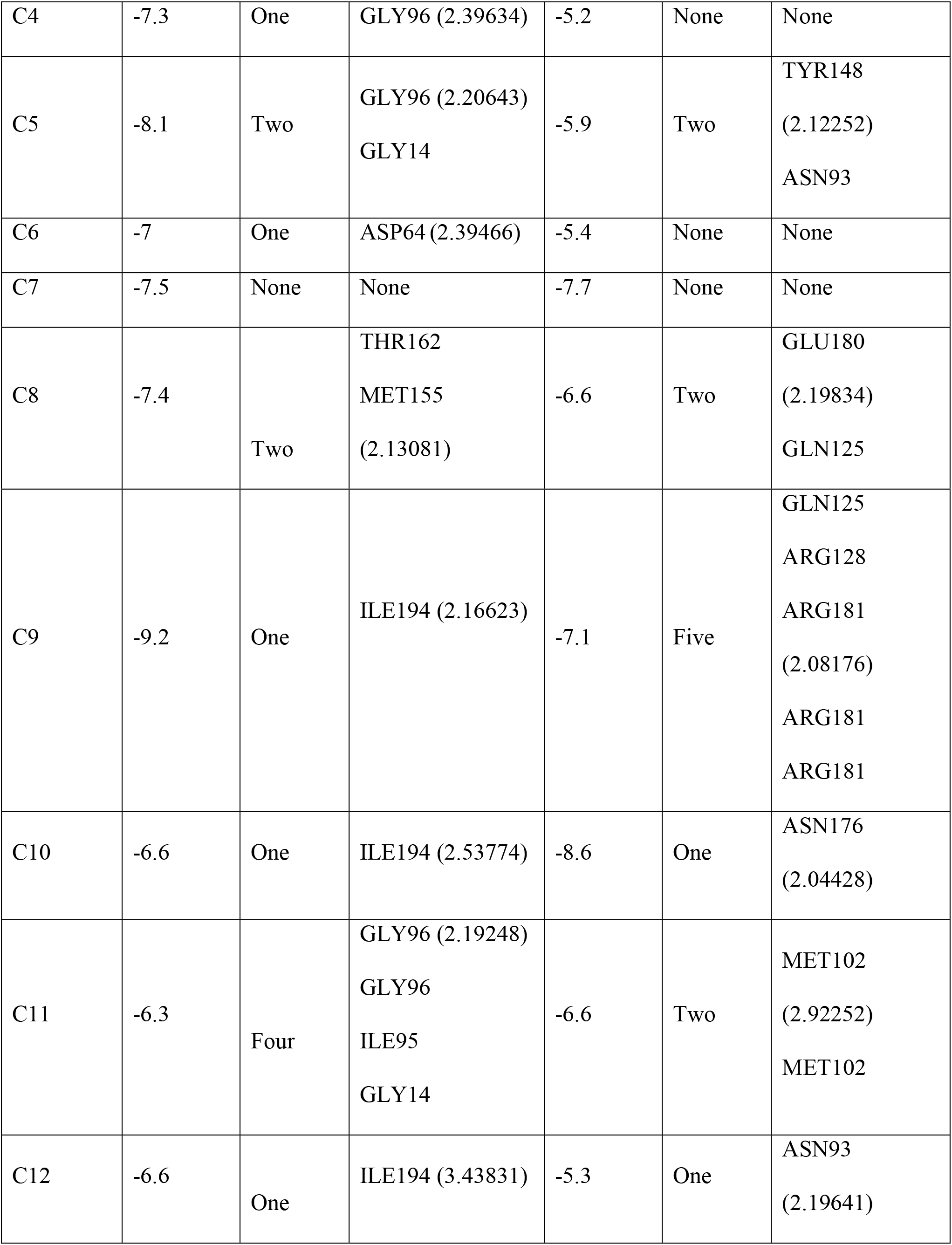
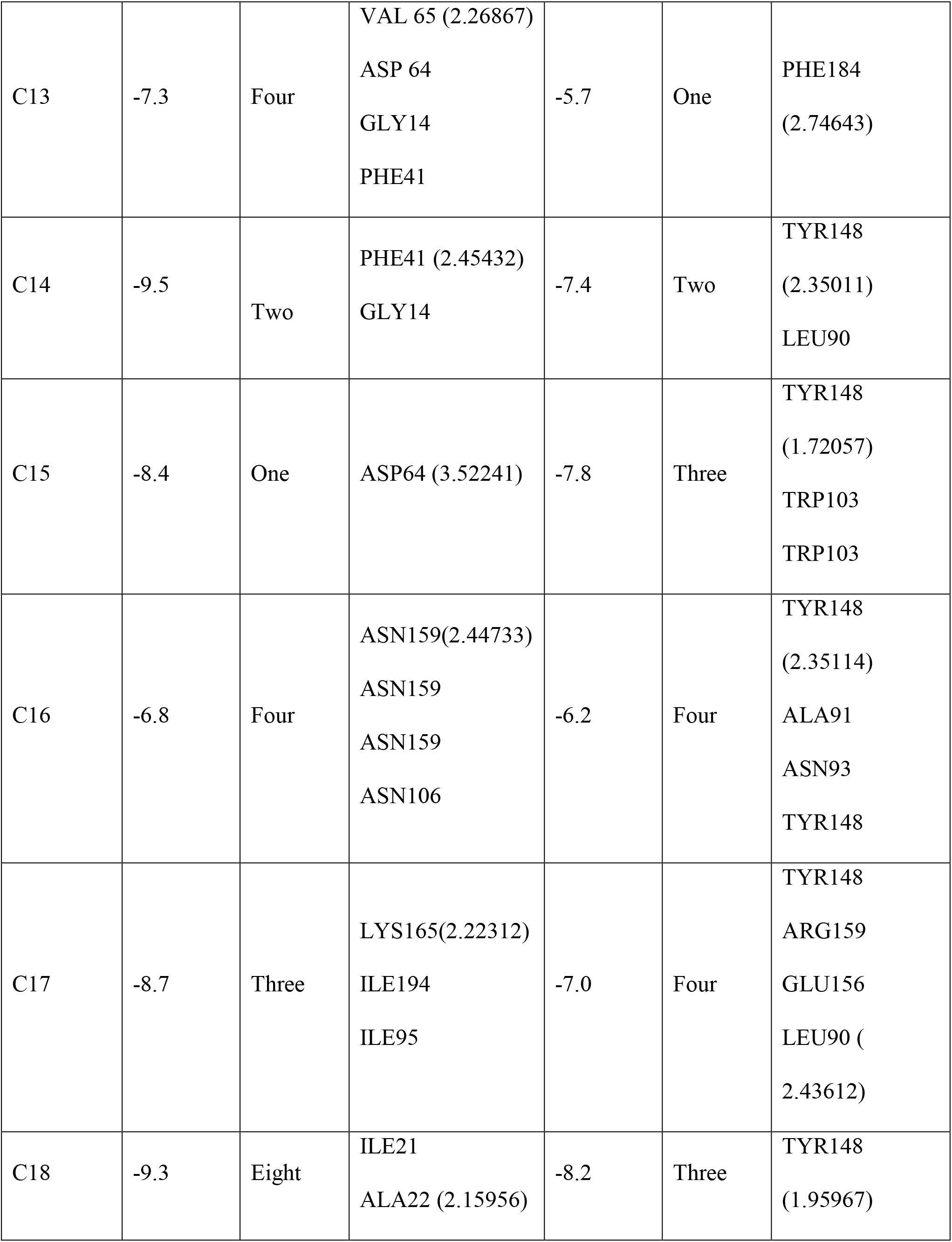
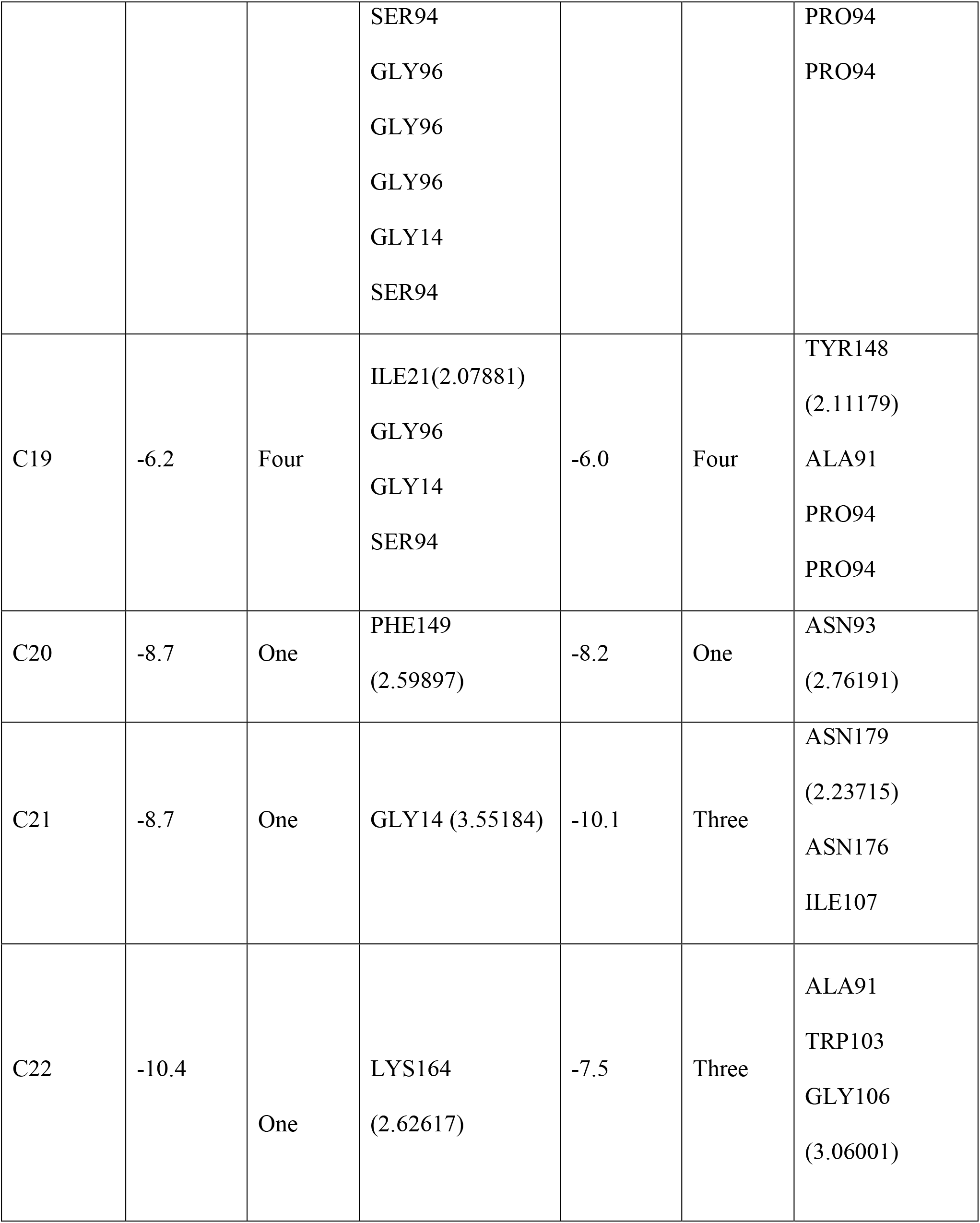
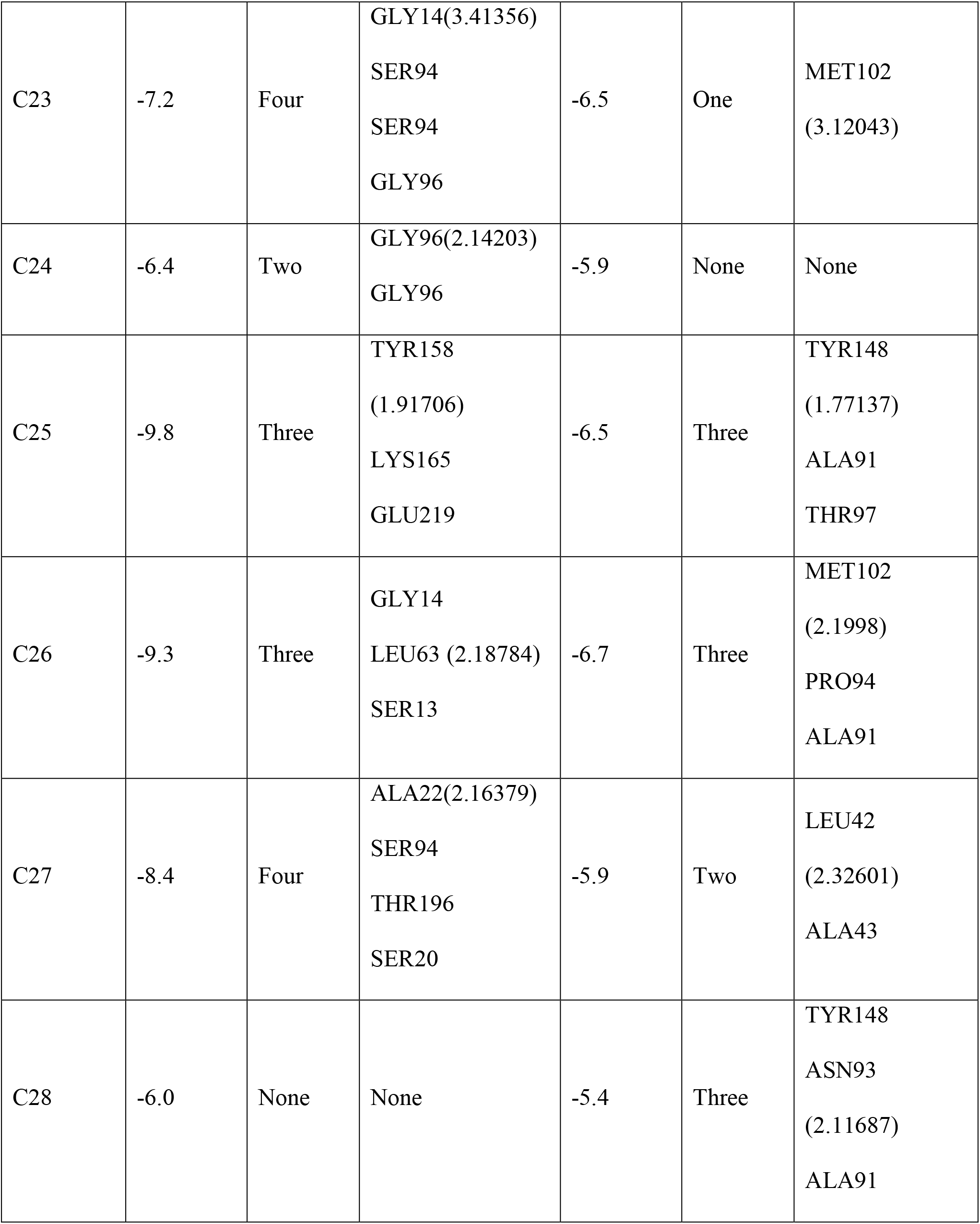
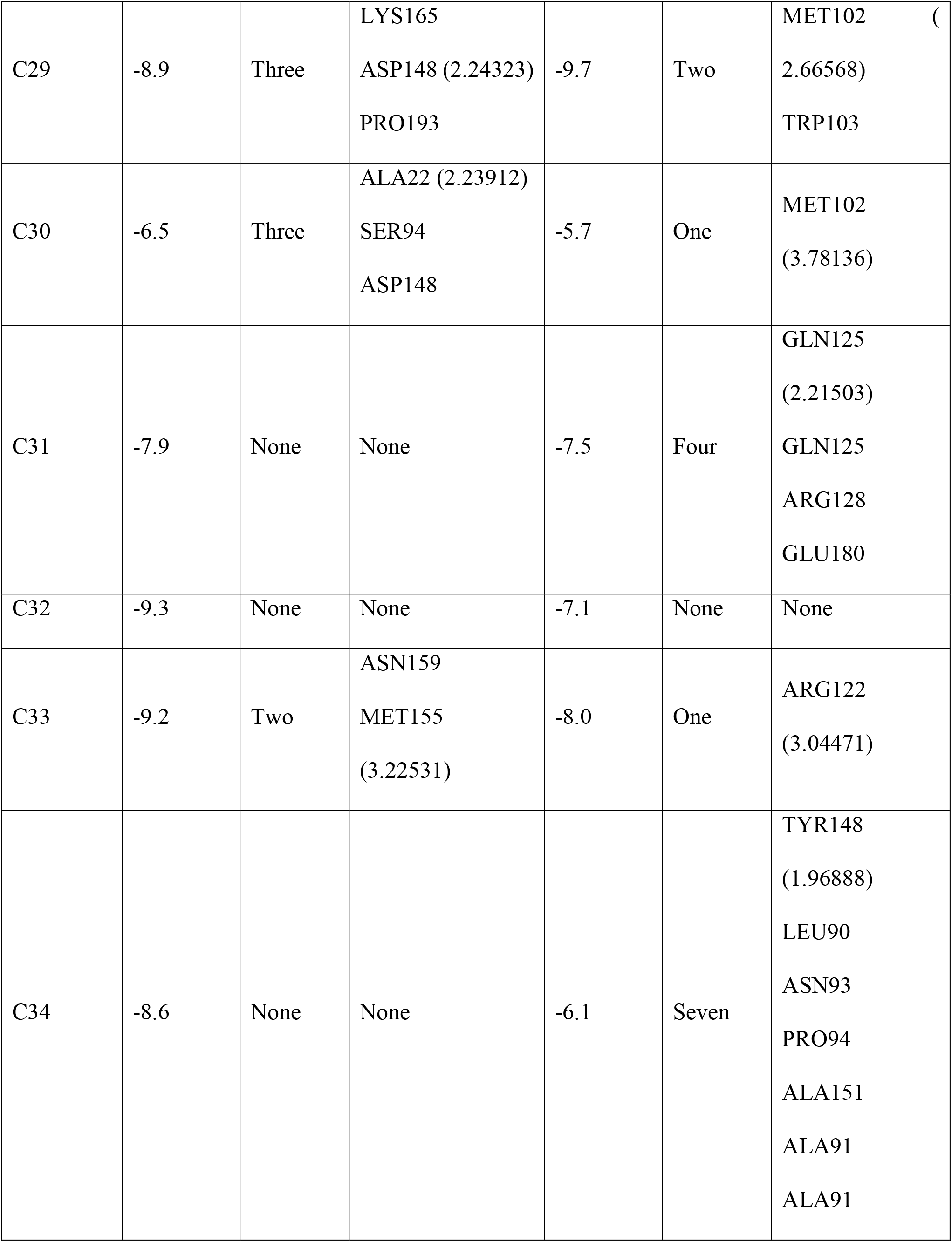
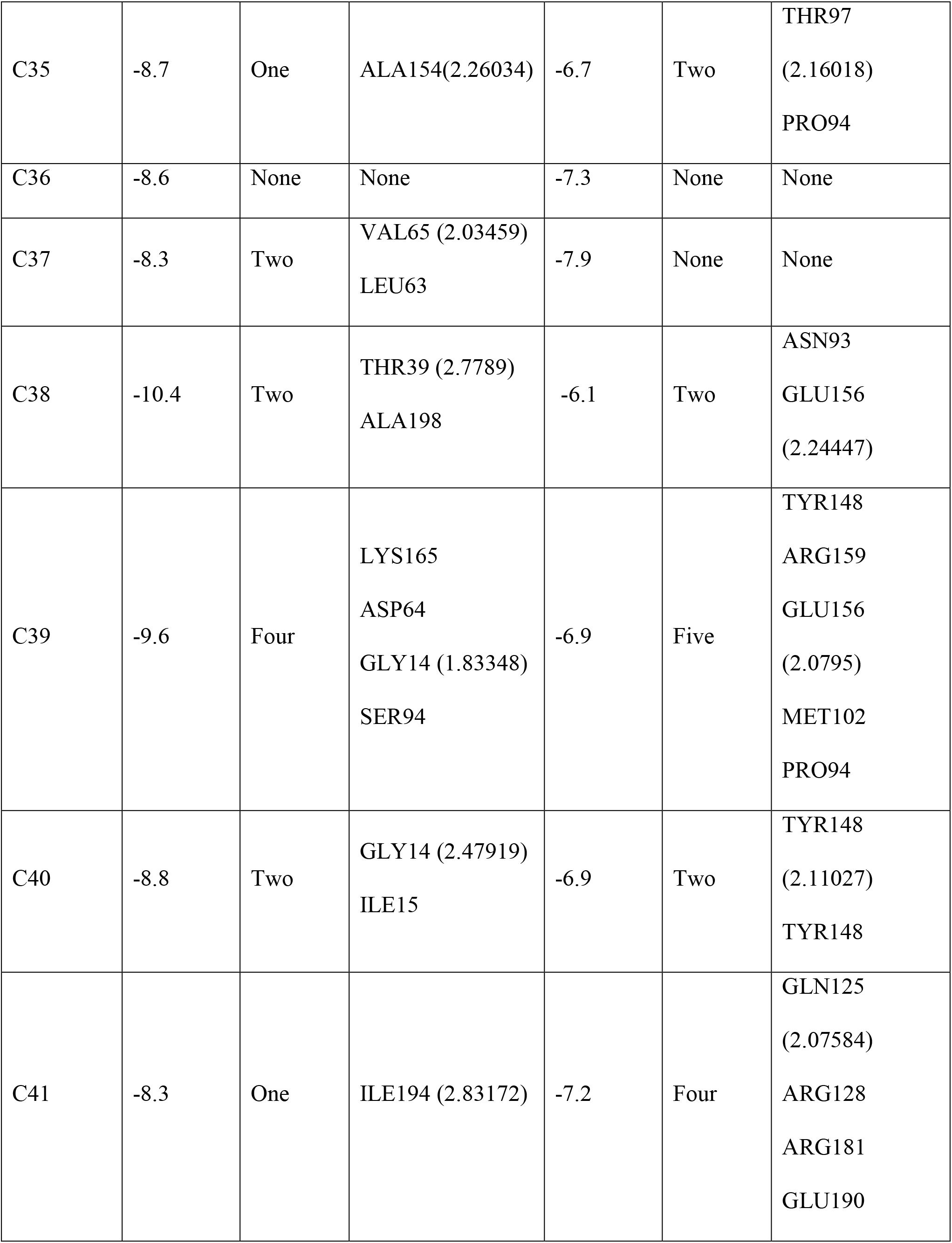
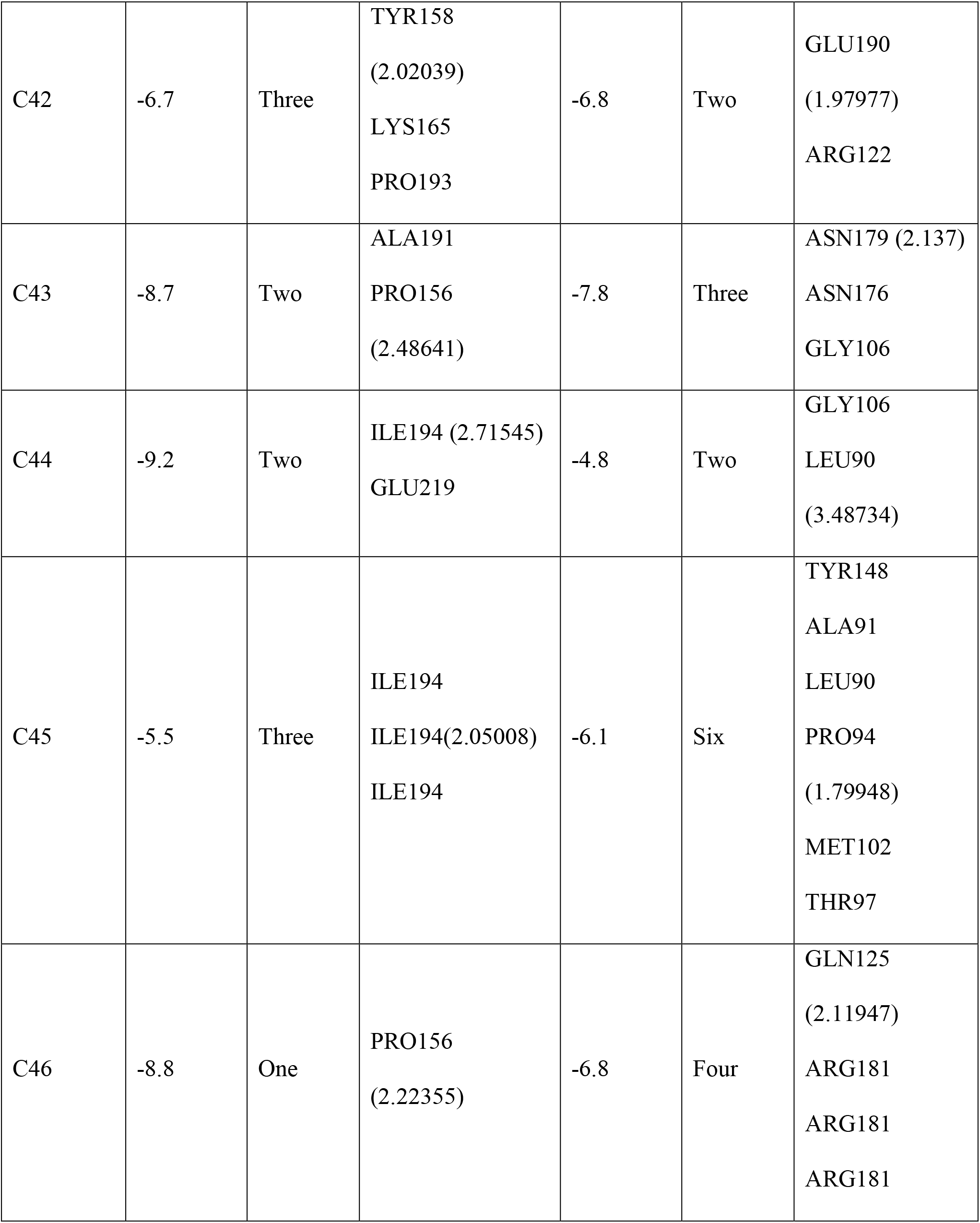
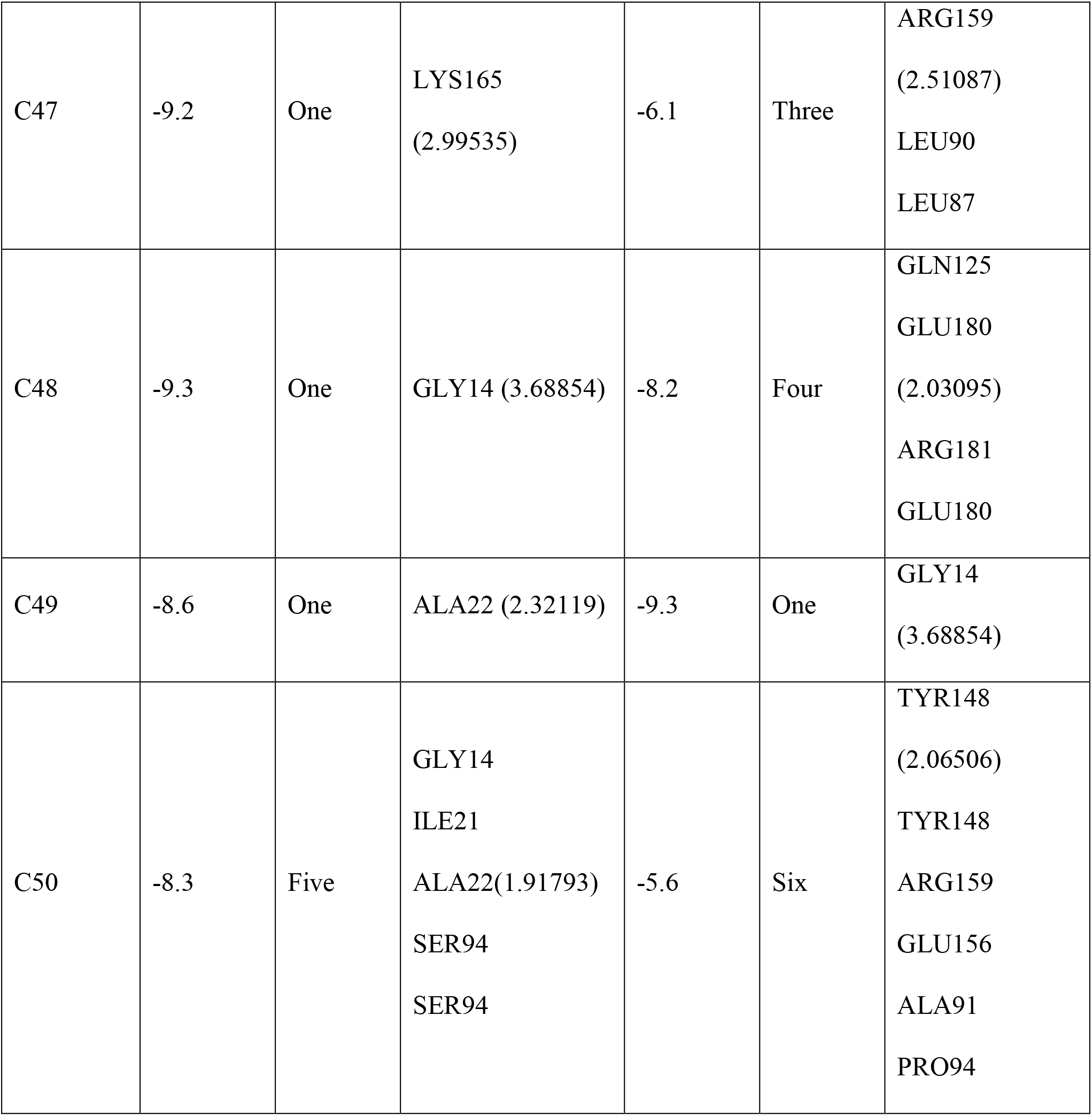
Molecular docking results and Hydrogen bond interaction between receptors (InhA, EthR) and fifty ligands, along with control drugs by Autodock Vina.

### 3.2. P450 Site of Metabolism (SOM) prediction

Cytochromes P450 plays an important role in metabolizing external materials like drugs. The probable metabolic sites of CYP (1A2, 2A6, 2B6, 2C8, 2C9, 2C19, 2D6, 2E1, 3A4 and combined) of the four ligands:-3-[3-(4-Fluorophenyl)-1,2,4-oxadiazol-5-yl]-N-(2-methylphenyl)piperidine-1-carboxamide (C22) ; Maritinone (C38) ; 3-Methyl-benzofuran-2-carboxylic acid pyridin-4-ylamide (C21) and 5-(4-Ethyl-phenyl)-2-(1H-tetrazol-5-ylmethyl)-2H-tetrazole (C29) were organized using RS-Web Predictor tool. The possible sites of metabolism by the isoforms were indicated by circles on the chemical structure of the four ligands. The probable sites of P450 metabolism are shown in Table 3.

**Table 3:**
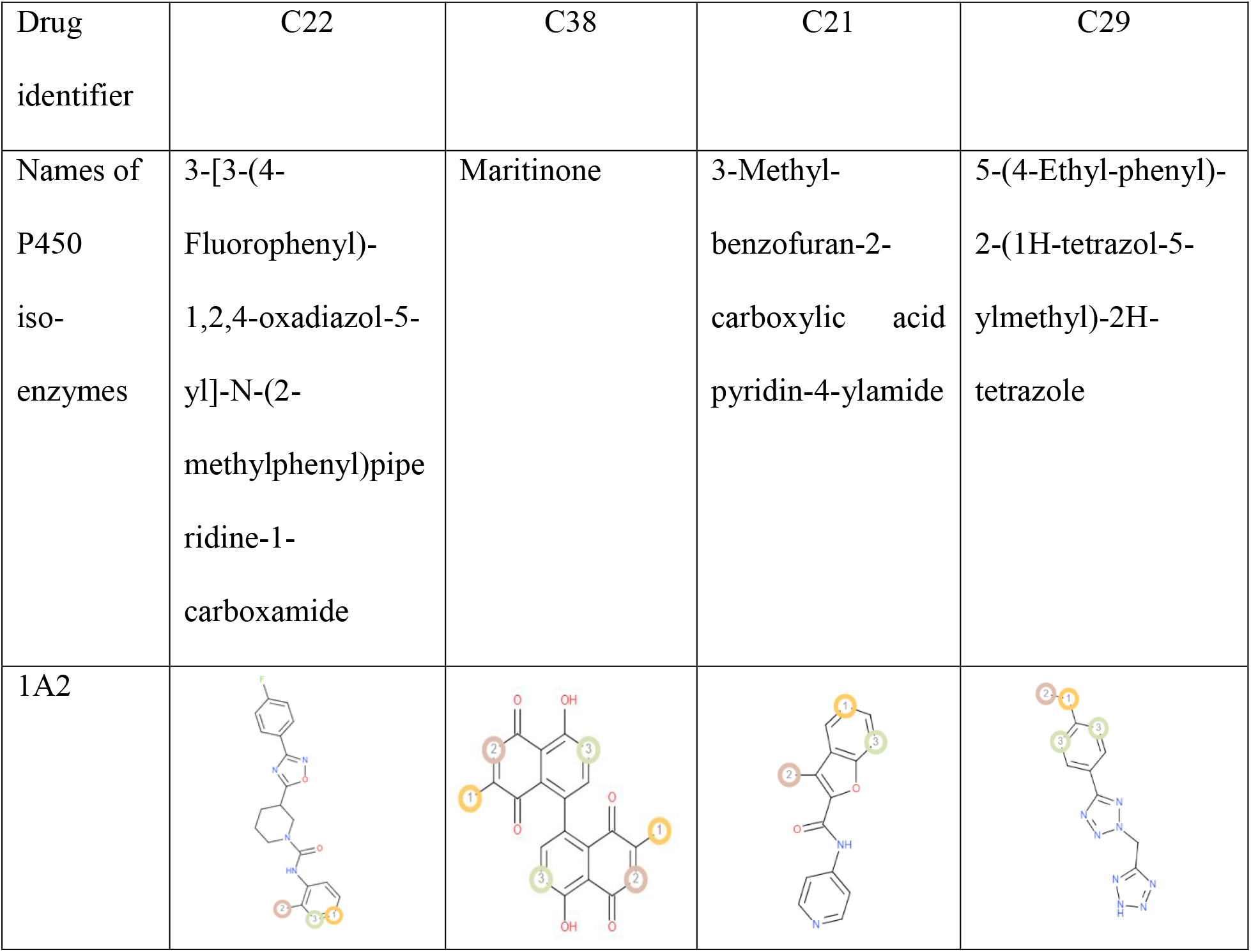
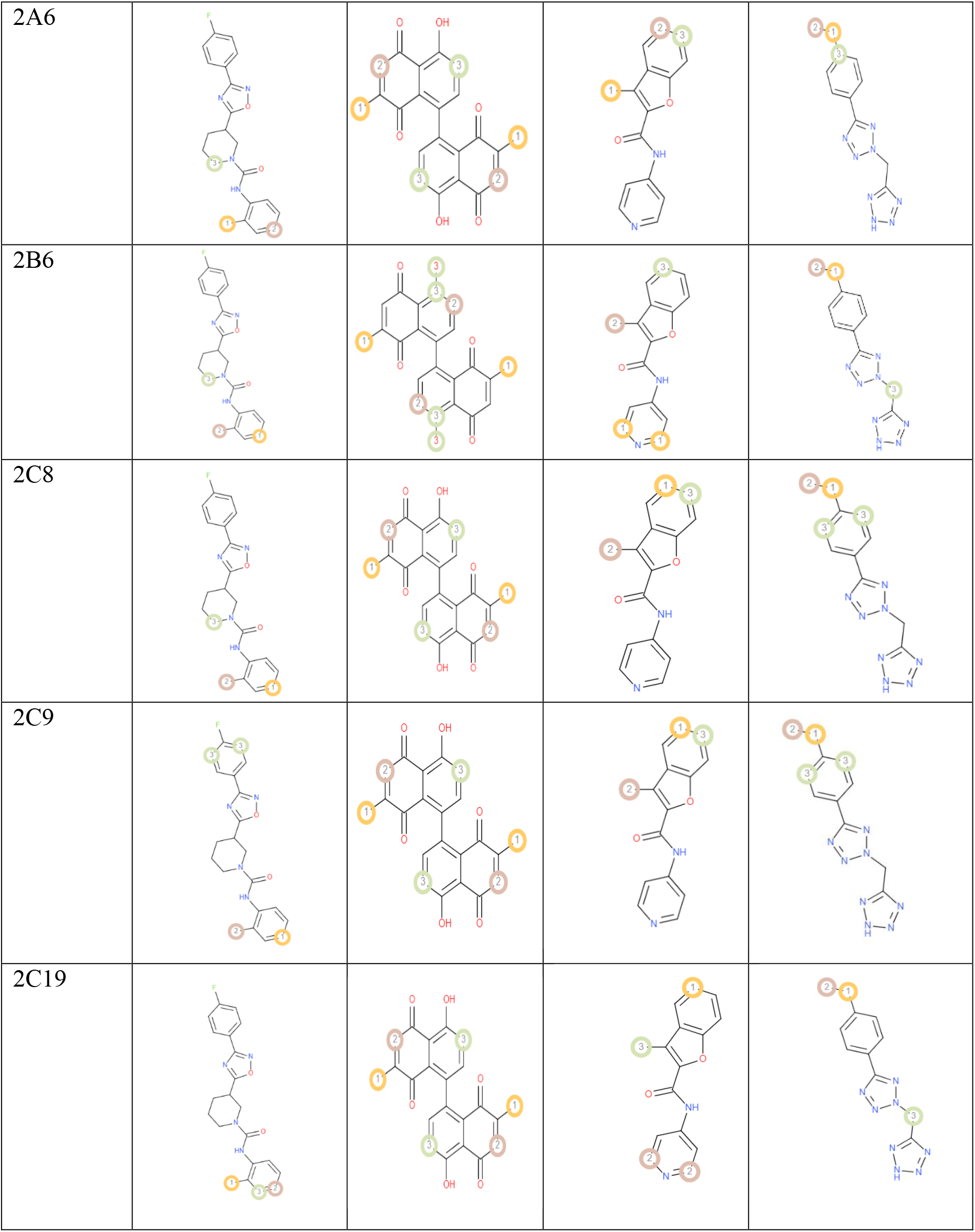
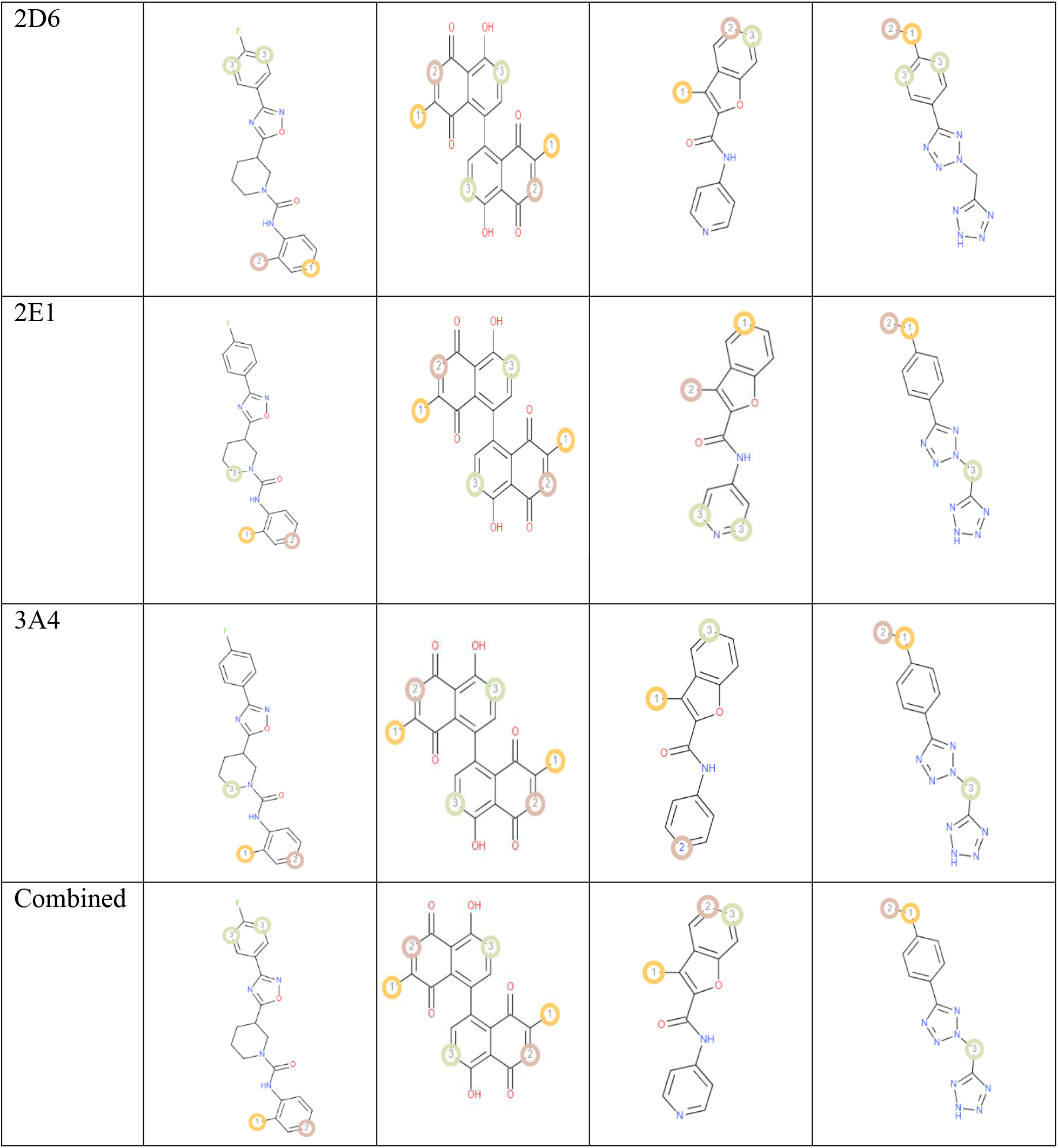
The P450 Site of Metabolism (SOM) prediction results of the best four ligands.

### 3.3. Drug-likeness and ADMET analysis

The concept of drug-likeness has appeared as an approach that can screen the high-affinity ligands with acceptable ADME (absorption, distribution, metabolism, excretion) properties. In our study, phytochemicals: 3-[3-(4-Fluorophenyl)-1,2,4-oxadiazol-5-yl]-N-(2-methylphenyl)piperidine-1-carboxamide (C22) ; Maritinone (C38) ; 3-Methyl-benzofuran-2-carboxylic acid pyridin-4-ylamide (C21) and 5-(4-Ethyl-phenyl)-2-(1H-tetrazol-5-ylmethyl)-2H-tetrazole (C29) were subjected to Drug-likeness and ADMET analysis and all of them followed the Lipinski’s rule of five criteria: molecular weight (acceptable range: <500), number of hydrogen bond donors (acceptable range: ≤5), number of hydrogen bond acceptors (acceptable range: ≤10), lipophilicity (expressed as LogP, acceptable range: <5) and molar refractivity (40-130). C38 had the highest topological polar surface area (108.74 Å^2^) and C21 had the lowest polar surface area (55.13 Å^2^) although all of the five compounds satisfy the ideal value (20-130 Å2). The Ghose, Veber, Egan, Muegge rules are followed by all of the four compounds. The number of rotatable bonds, bioavailability scores, log S fell within the standard range for the four compounds (Table 4).

**Table 4:**
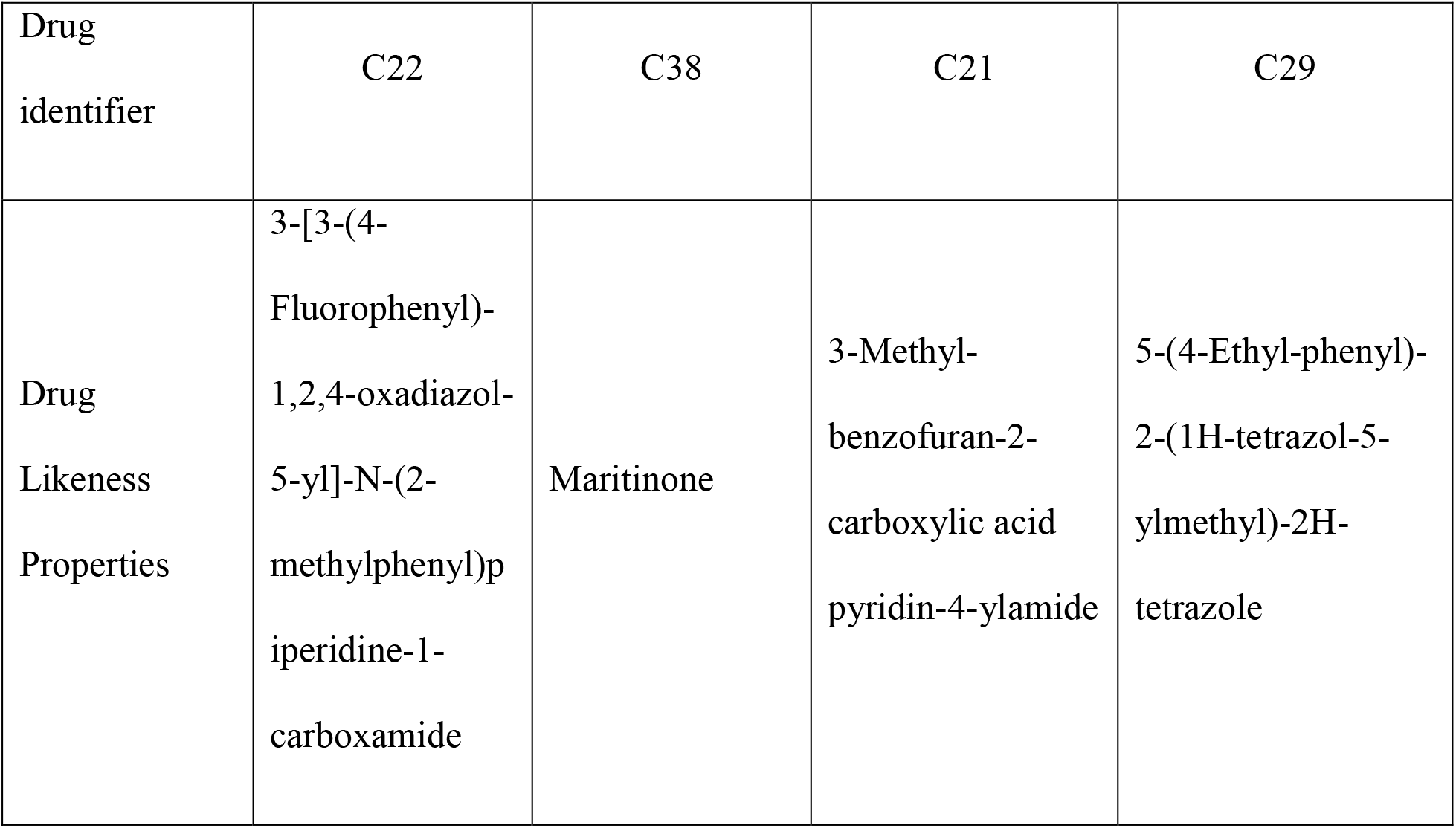
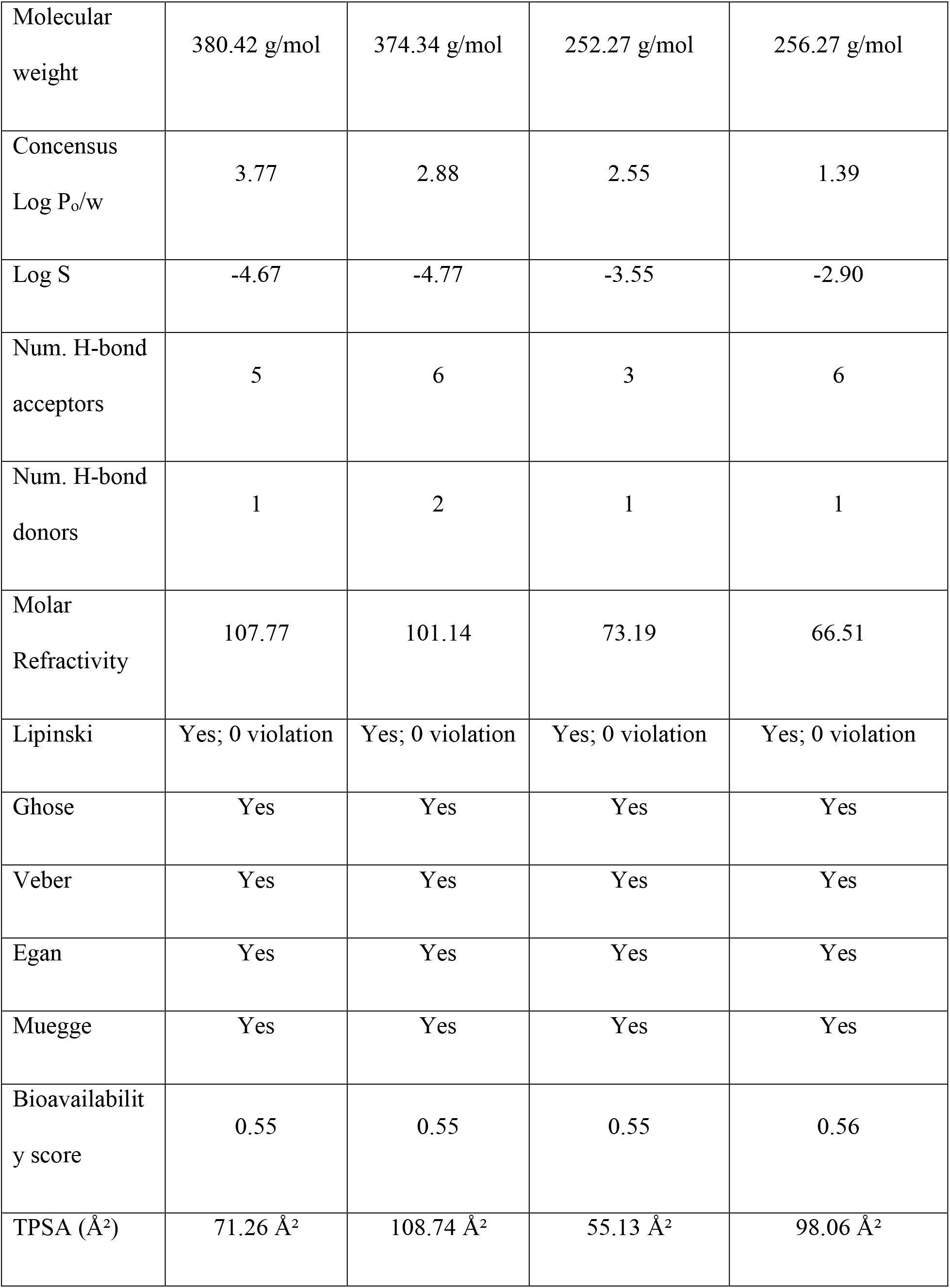
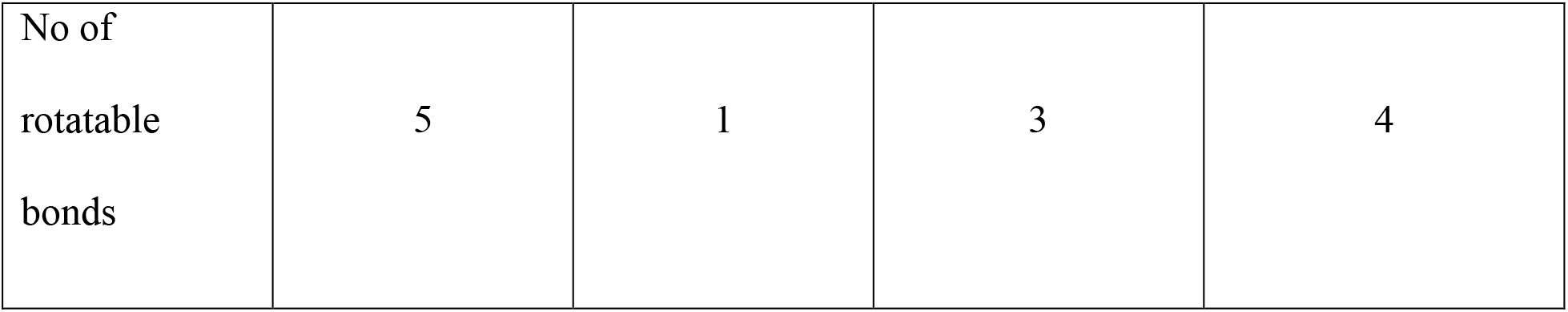
The Drug-Likeness properties of the best four ligands.

The relative ADMET profiles of screened ligands are described in Table 5. The evaluated pharmacokinetic data showed that all of the selected molecules had a high intestinal absorption rate and oral bioavailability. Each chosen molecule was capable of pervading Caco2 cell lines. C22 acted as P-glycoprotein substrate, P-glycoprotein inhibitor and substrate for CYP3A4 cytochrome.

**Table 5:** ADMET Prediction of the best four ligands.

| Drug identifier | C22 | C38 | C21 | C29 |
| --- | --- | --- | --- | --- |
| Properties | 3-[3-(4-Fluorophenyl)-1,2,4-oxadiazol-5-yl]-N-(2-methylphenyl)piperidine-1-carboxamide | Maritinone | 3-Methyl-benzofuran-2-carboxylic acid pyridin-4-ylamide | 5-(4-Ethylphenyl)-2-(1H-tetrazol-5-ylmethyl)-2H-tetrazole |
| Human Intestinal Absorption | Positive (0.9907) | Positive (0.9952) | Positive (0.9726) | Positive (0.9933) |
| Blood Brain Barrier | Positive<br>(0.9871) | Negative<br>(0.7104) | Positive<br>(0.9880) | Positive<br>(0.9825) |
| Caco-2 | Positive<br>(0.5586) | Positive<br>(0.6857) | Positive<br>(0.8079) | Positive<br>(0.7131) |
| Human oral<br>bioavailability | Positive<br>(0.5857) | Positive<br>(0.6286) | Positive<br>(0.8429) | Positive<br>(0.6000) |
| Subcellular<br>localization | Mitochondria<br>(0.7027) | Mitochondria<br>(0.9018) | Mitochondria<br>(0.4417) | Mitochondria<br>(0.8104) |
| P-glycoprotein<br>inhibitor | Positive<br>(0.6537) | Negative<br>(0.8316) | Negative<br>(0.8282) | Negative<br>(0.9441) |
| P-glycoprotein<br>substrate | Positive<br>(0.6480) | Negative<br>(0.9566) | Negative<br>(0.8627) | Negative<br>(0.5591) |
| CYP3A4 substrate | Positive<br>(0.6532) | Negative<br>(0.5992) | Positive<br>(0.5080) | Negative<br>(0.5554) |
| CYP2C9 substrate | Positive<br>(0.5894) | Negative<br>(0.8009) | Negative<br>(1.0000) | Negative<br>(0.7957) |
| CYP2D6<br>substrate | Negative<br>(0.8300) | Negative<br>(0.8664) | Negative<br>(0.8972) | Negative<br>(0.8707) |
| CYP3A4 inhibition | Negative<br>(0.5188) | Negative<br>(0.7006) | Positive<br>(0.7887) | Negative<br>(0.9575) |
| CYP2C9 inhibition | Positive<br>(0.5217) | Positive<br>(0.9592) | Positive<br>(0.7423) | Negative<br>(0.8643) |
| CYP2C19 inhibition | Negative<br>(0.5209) | Positive<br>(0.7766) | Positive<br>(0.8633) | Negative<br>(0.6388) |
| CYP2D6 inhibition | Negative<br>(0.9132) | Negative<br>(0.7041) | Negative<br>(0.5689) | Negative<br>(0.9152) |
| CYP1A2 inhibition | Negative<br>(0.7245) | Positive<br>(0.9159) | Positive<br>(0.9725) | Positive<br>(0.7147) |
| Hepatotoxicity | Negative<br>(0.5750) | Positive<br>(0.9250) | Positive<br>(0.9500) | Negative<br>(0.5000) |
| Carcinogenicity<br>(binary) | Negative<br>(0.8857) | Negative<br>(0.6510) | Negative<br>(0.8571) | Negative<br>(0.9143) |

Neither of the ligands acted as CYP2D6 substrate nor CYP2D6 inhibitor. Only C29 showed an inhibitory effect on CYP2C9 cytochrome. C22 acted as inhibitor for CYP3A4, CYP2C19, CYP1A2 cytochrome but acted as a substrate for CYP2C9 cytochrome. C21 inhibited CYP3A4, CYP2C19, CYP1A2 cytochrome isoform but represented as a substrate for CYP3A4 isoform.

All selected ligands were non-carcinogenic. C38 and C21 were both hepatotoxic. Overall, considering the scores 3-[3-(4-Fluorophenyl)-1,2,4-oxadiazol-5-yl]-N-(2-methylphenyl) piperidine-1-carboxamide (C22) and 5-(4-Ethyl-phenyl)-2-(1H-tetrazol-5-ylmethyl)-2H-tetrazole (C29) ligand went along with the maximum criteria. The best ligands and control drugs with respective receptors (C22 and InhA complex, Isoniazid and InhA complex, C29 and EthR complex, Ethionamide and EthR complex) were depicted in Figure (3 & 4).

**Figure 3:**
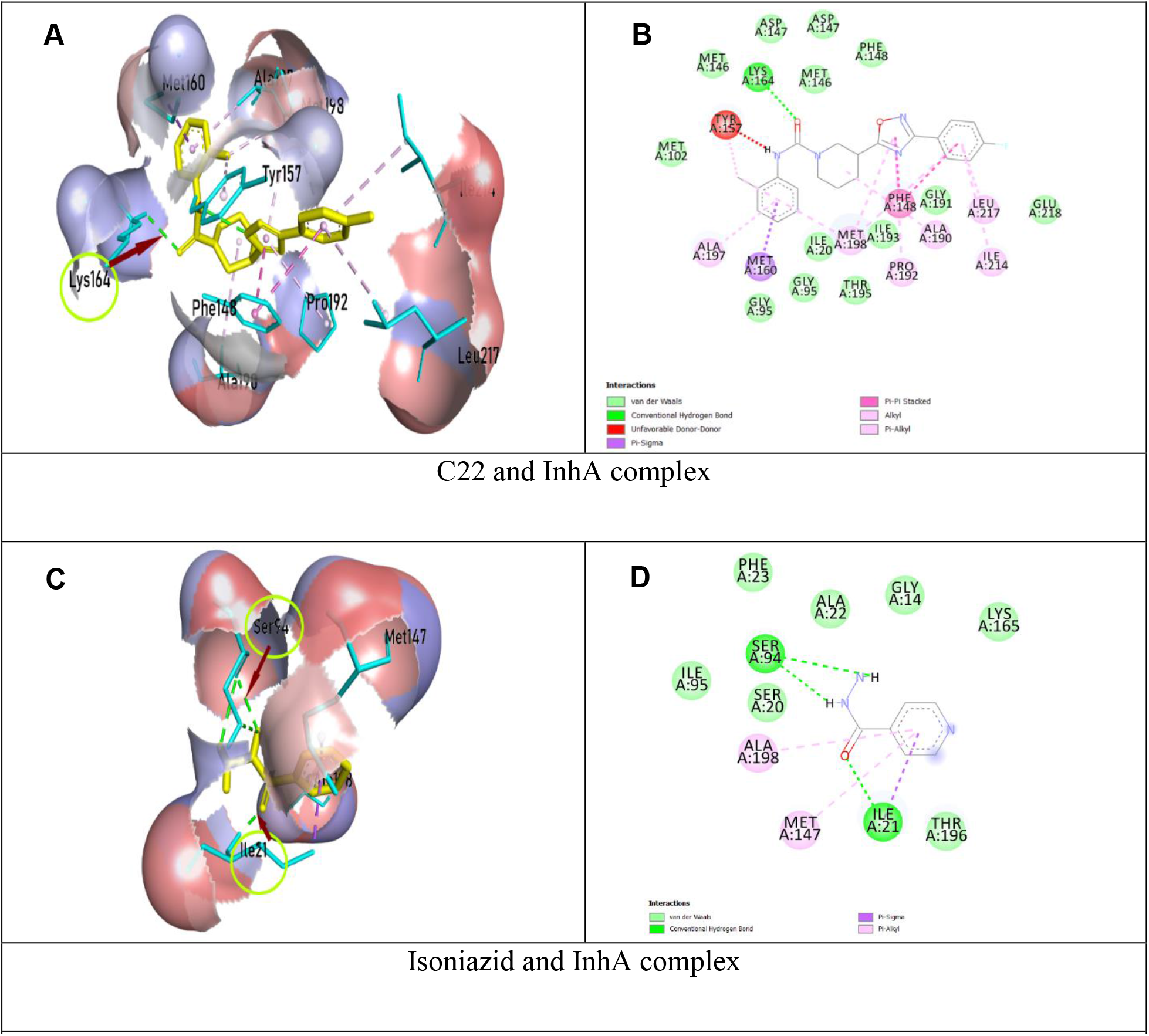
Schematic representation of InhA and drug (C22, Isoniazid) complex; On the left, ligands were in yellow color, parts of protein in cyan color, hydrogen bonds and interacting amino acids were shown with arrows and circle; On the right, two-dimensional image of InhA and drug (C22, Isoniazid) interaction were shown, green dotted line denoted hydrogen bonds.

**Figure 4:**
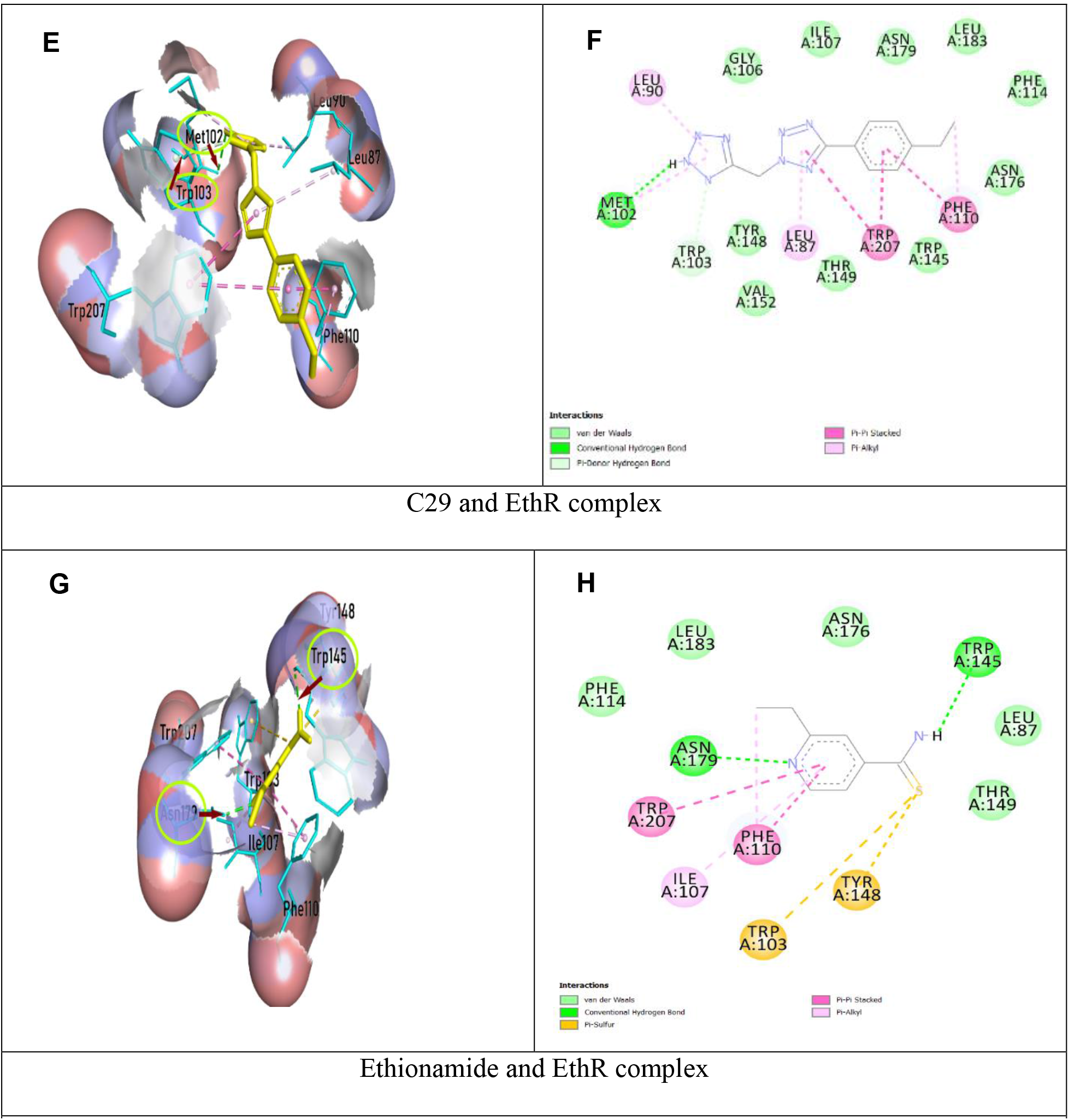
Schematic representation of EthR and drug (C29, Ethionamide) complex; On the left, ligands were in yellow color, parts of protein in cyan color, hydrogen bonds and interacting amino acids were shown with arrows and circle; On the right, two-dimensional image of EthR and drug (C29, Ethionamide) interaction were shown, green dotted line denoted hydrogen bonds.

### 3.4. Molecular Dynamics Simulation

In our study, Molecular Dynamics Study was performed to assess conformational stability of protein-ligand complexes as well as InhA and EthR receptors. The overall conformational similarity between control drugs and the drugs being tested with target proteins was compared assessing Root Mean Square deviation (RMSD). As represented in Figure 5(A), RMSD of Free InhA protein continued stability between 1 ns to 5 ns timescale at about 1.4 Å and 5 ns (nano second) to 9 ns at about 1.7 Å. (Isoniazid +InhA) complex displayed steadiness in RMSD between 1 ns to 5 ns timescale at around 1.5 Å and 8 ns to till the end of run at around 1.8 Å. (C22+InhA) complex held stable backbone stability from 3 ns to 7 ns, with RMSD around 2 Å. Free EthR protein, (Ethionamide+EthR) protein-ligand complex and (C29+EthR) complex showed a backbone RMSD under 2 Å, which suggested minor structural changes. Free EthR protein remained stable between 1 ns to 6 ns, exhibiting RMSD around 1.2 Å. (Ethionamide+EthR) complex also showed stability from 1 ns to 6 ns timeline, with a RMSD backbone of 1.2 Å, then exhibiting small increased RMSD. On the other hand, RMSD of (C29+EthR) complex persisted stable from 1 ns to 5 ns timeline at about 1.5 Å, then after slight fluctuation, remained stable from 8 ns to the end of run on average 1.4 Å.

**Figure 5:**
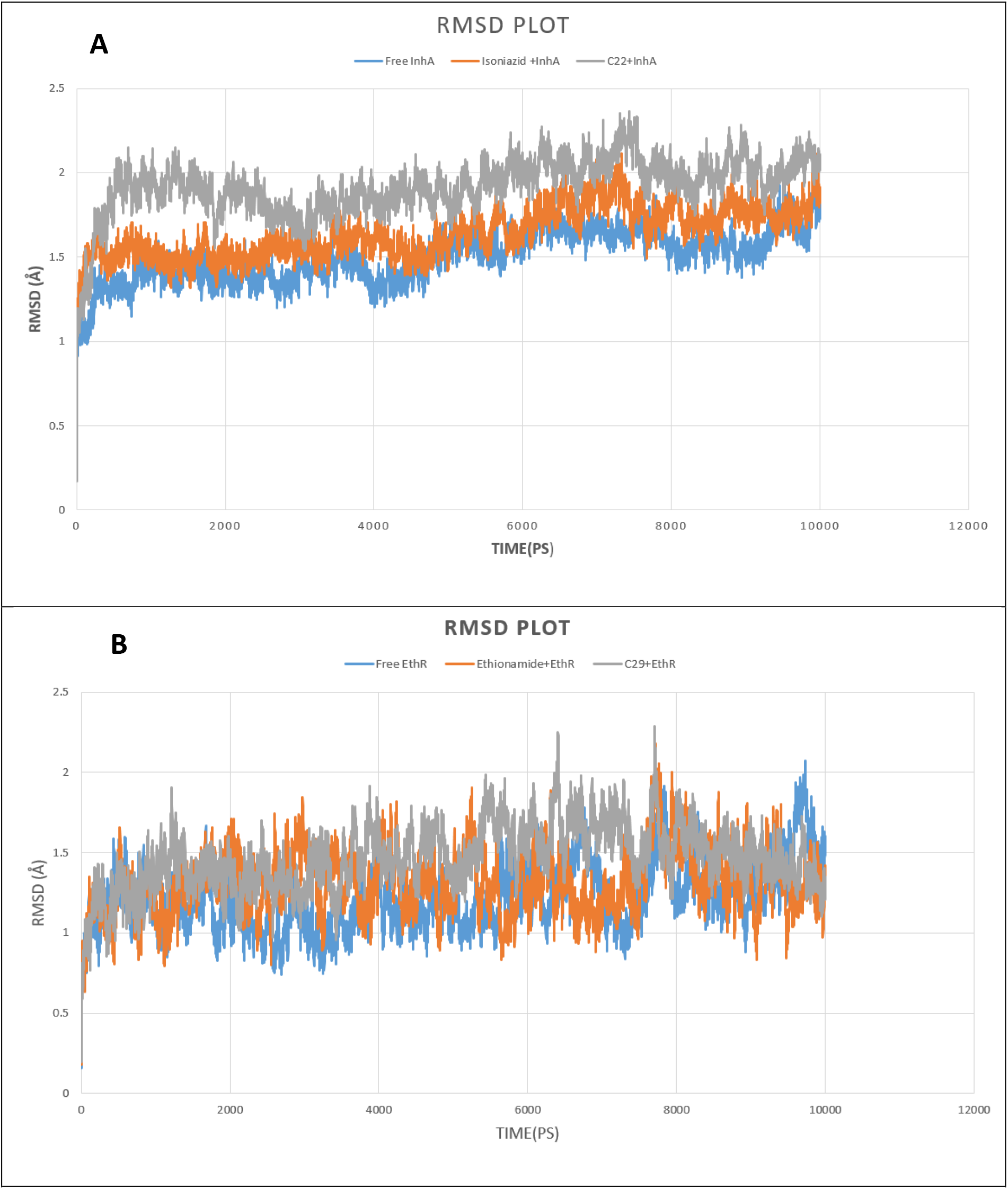
10 ns Molecular Dynamics Simulation (MDS) RMSD of Free protein and bounded protein; (A, B) Free protein (InhA and EthR) in blue color, Control drugs (Isoniazid and Ethionamide) and protein in orange color, Selected drug (C22 + C29) and protein in grey color.

Root Mean Square Fluctuation (RMSF) analysis per residue for backbone atoms was conducted to assess changes in conformation of Cα backbone of the systems. From figure 6(C), Free InhA, (Isoniazid +InhA), (C22+InhA) complexes mostly had RMSF from 0.4 to 2 Å that indicated close conformational contact between protein and ligands. Nevertheless, the higher fluctuation of RMSF between 205 to 211 residues confirmed the presence of loop within this region. Our MDS study showed that the RMSF of Free EthR, (EthR+Ethionamide), (EthR+C29) complexes fluctuated mostly between 0.5 to 1.5 Å, indicating close contact between active pocket of receptors and drugs. However, higher fluctuation from 73 to 75 and 170 to 172 amino acid residues indicated that the free protein and its complexes were within the loop regions.

**Figure 6:**
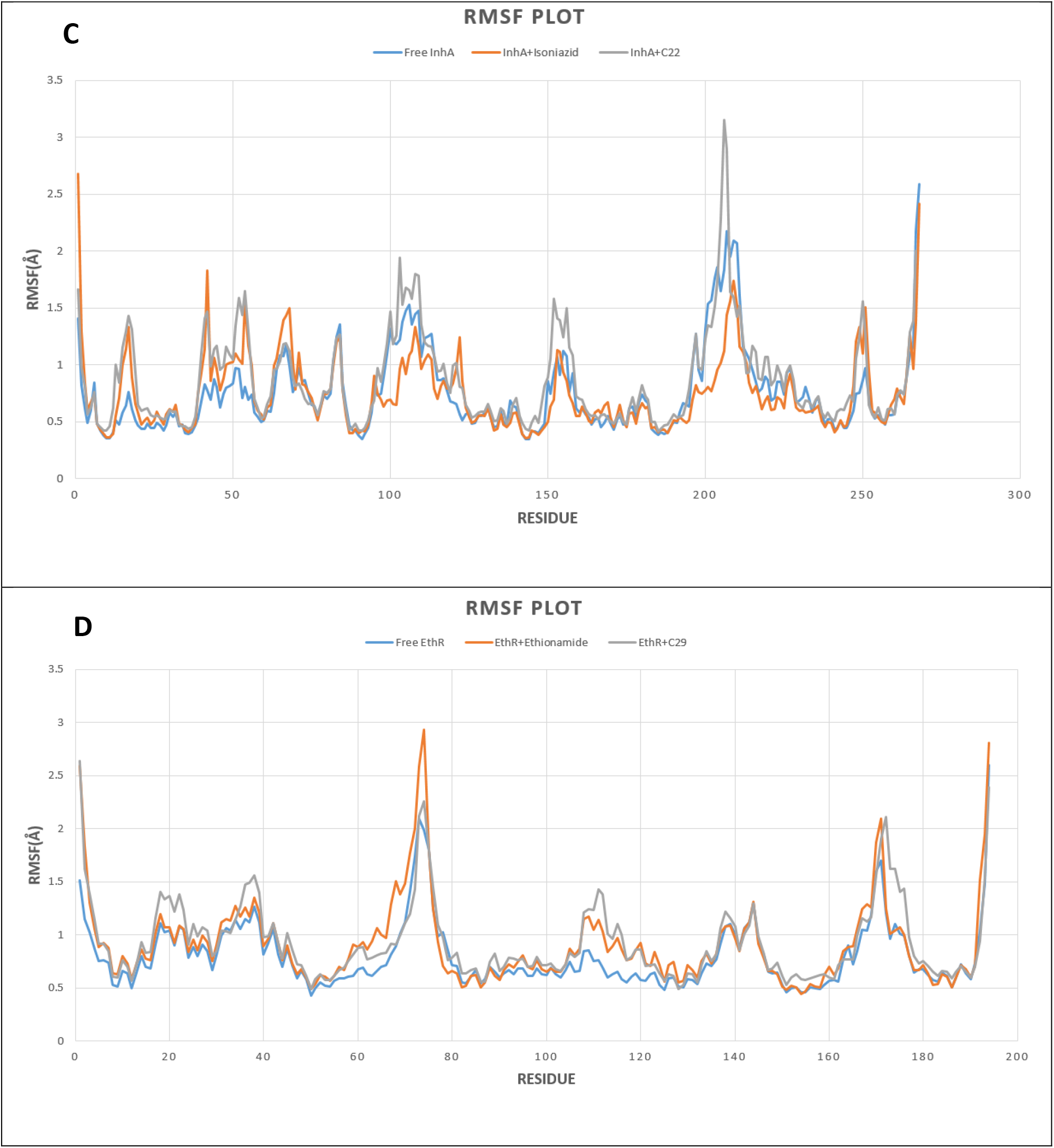
RMSF outline of Free protein and bounded protein; (C, D) Free protein (InhA and EthR) in blue color, Control drugs (Isoniazid and Ethionamide) and protein in orange color, Selected drug (C22 + C29) and protein in grey color.

Radius of Gyration with time was calculated to assess the change of compactness after ligand binding with receptors. From figure 7(E), the Rg of Free InhA was reported between 17.9 to 18.2 Å; (InhA+Isoniazid) complex showed Rg between 17.9 to 18.3 Å; (InhA+C22) complex continue Rg between 18.1 to 18.4 Å through the simulation. This evaluation proved ligand binding did not affect the compactness of the protein. On the other hand, from figure 7(F), it was evident Free EthR had stable Rg value around 19.3 Å from 2 to 8 ns; (EthR+Ethionamide) complex remained stable Rg about 19.4 Å between 5 ns to till the end of simulation and (EthR+C29) complex kept stability from 5 to 9 ns, with Rg around 19.3 Å. This confirmed ligand binding with respective receptor did not cause structural instability to proteins.

**Figure 7:**
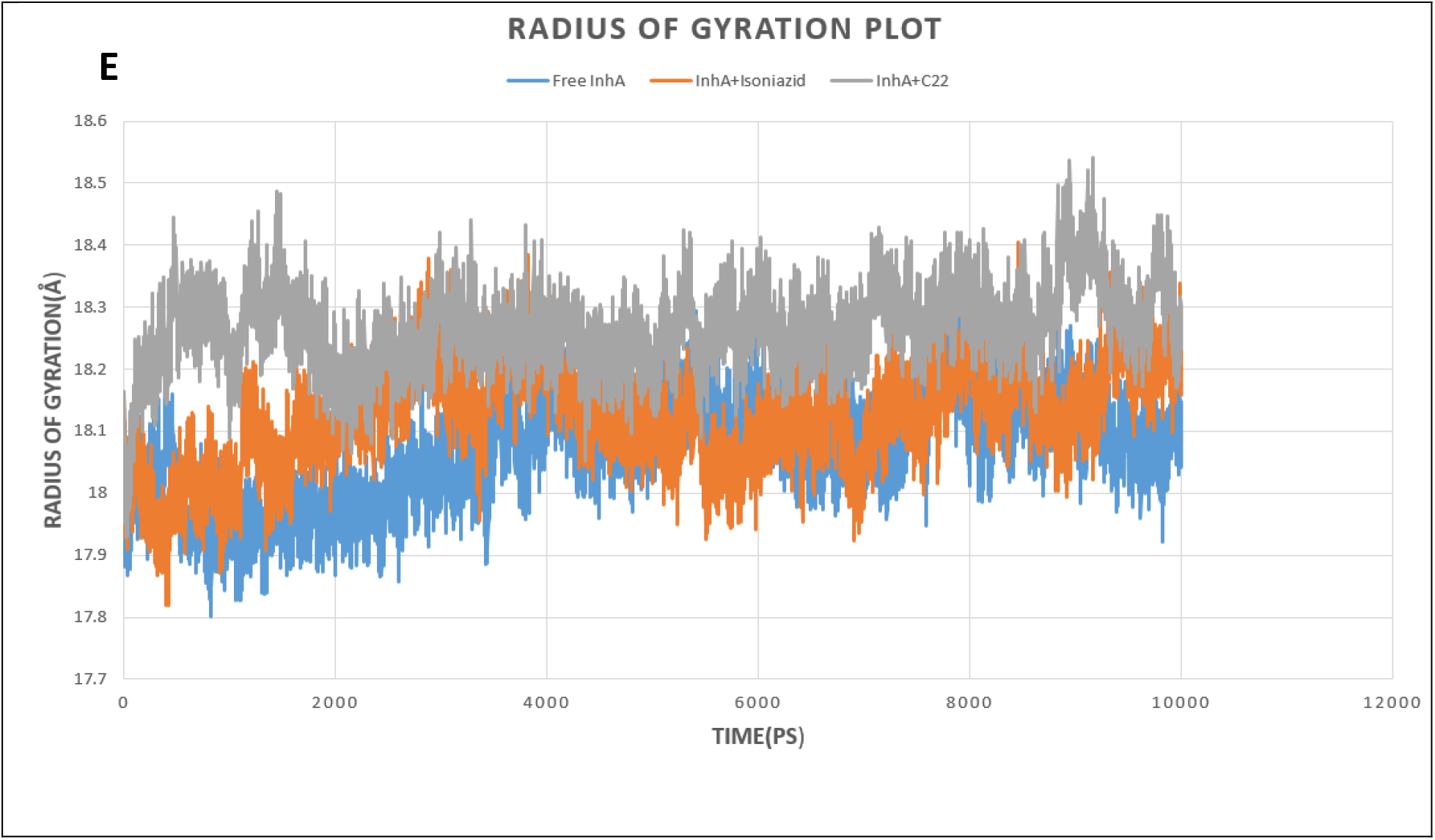
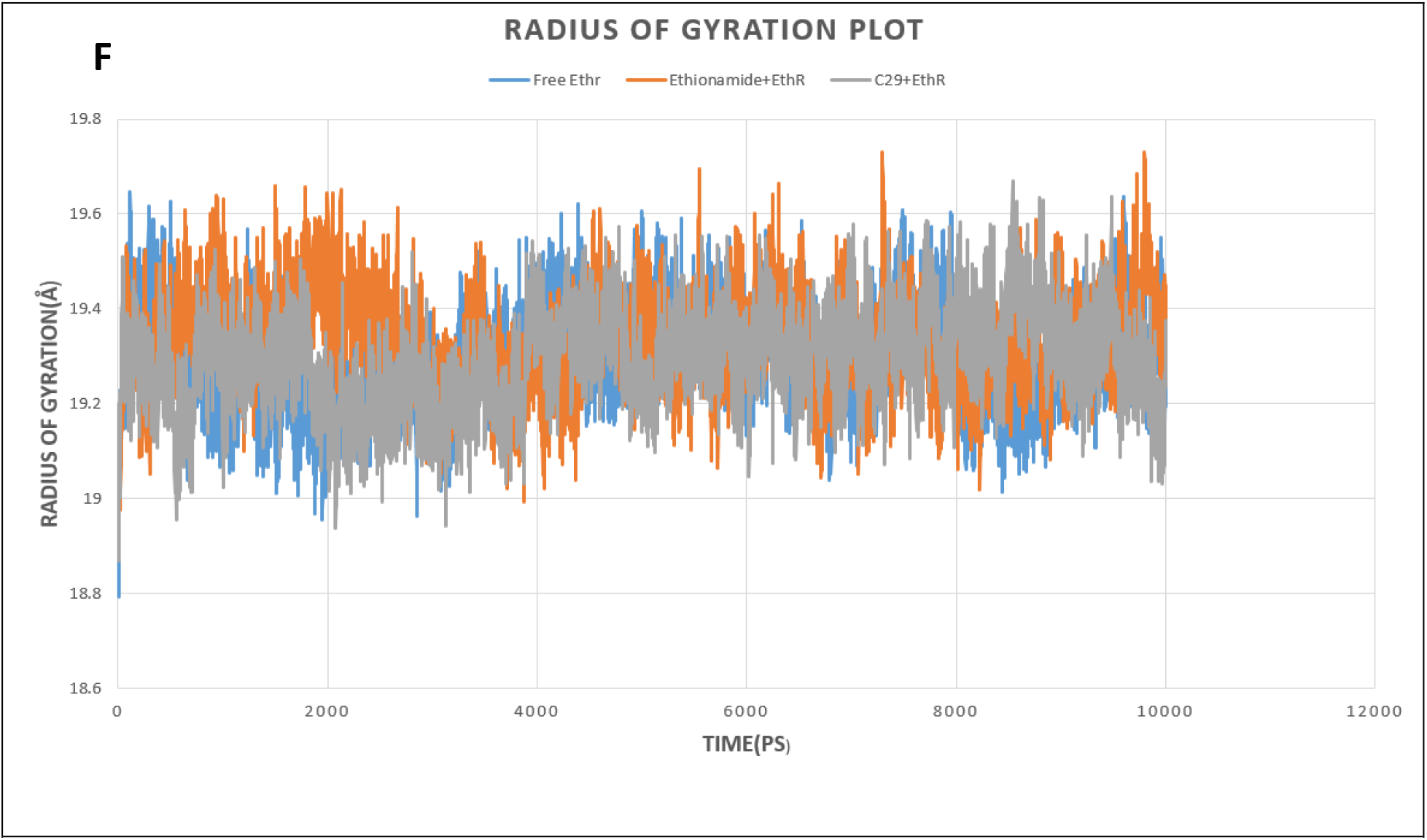
Analysis of Radius of Gyration (Rg); (E, F) Free protein (InhA and EthR) in blue color, Control drugs (Isoniazid and Ethionamide) and protein in orange color, Selected drug (C22 + C29) and protein in grey color.

## 4. DISCUSSION

In our study, the screening of high affinity ligands was accomplished by two stages of molecular docking analysis. As an initial docking evaluation, fifty ligands were primarily docked separately against the InhA and EthR receptors protein via the DockThor server [29]. Both for InhA and EthR proteins, all the compounds showed lower affinity score than their respective controls. Subsequently Autodock Vina software was employed for computing binding energy and bound conformation in which two ligands for InhA protein and two ligands for EthR protein were chosen considering their lower binding score [32].

Drug-likeness is an approach that qualitatively evaluate the probability for a molecule to qualify as an oral drug with respect to bioavailability [46,47]. To analyze the drug-likeness property of our compounds we used SwissADME, a free web-based tool [40]. Lipinski’s Rule of 5, a rule of thumb was established to evaluate ‘drugability’ of new chemical entities having certain pharmacological or biological activity [48]. In the drug discovery and optimization approach, the Rule of 5 reveals that a drug compound should follow the criteria with the acceptable range having molecular weight: ≤500, number of hydrogen bond donors: ≤5, number of hydrogen bond acceptors: ≤10, lipophilicity (denoted as LogP): ≤5 and molar refractivity between 40 to 130 [36]. Our selected four compounds: 3-[3-(4-Fluorophenyl)-1,2,4-oxadiazol-5-yl]-N-(2-methylphenyl)piperidine-1-carboxamide (C22) ; Maritinone (C38) ; 3-Methyl-benzofuran-2-carboxylic acid pyridin-4-ylamide (C21) and 5-(4-Ethyl-phenyl)-2-(1H-tetrazol-5-ylmethyl)-2H-tetrazole (C29) followed the Lipinski’s rule of five criteria without violating any of its parameters. According to Ghose filter, to qualify as a drug, it should have log P value between −0.4 and 5.6, molecular weight from 160 to 480, molar refractivity between 40 and 130, total atomic count between 20 and 70 [37]. Veber rule states that compounds which meet two criteria of: 10 or fewer rotatable bonds and polar surface area equal or less than 140 Å are predicted to have good oral bioavailability [38]. The Egan rule considers candidate drug to have good oral bioavailability with −1.0 ≤logP ≤5.8 and TPSA ≤130 Å2 [49]. Muegge rule utilizes pharmacophore point filter to distinguish between drug-like and nondrug-like compounds which is based on simple structural rules. To pass the filtering, candidate drug should obtain two to seven pharmacophore points [39]. In addition to following the Lipiniski rule, they followed the Ghose, Veber, Egan and Muegge rules as well. The aqueous solubility of a compound expressed as log S determines the oral bioavailability resulting in affecting its absorption and distribution in the ADME/T properties [50].

The “Topological Polar Surface Area” (TPSA) is an approach that enables the calculation of the Polar Surface Area which is correlated with passive molecular transport through membranes, thus used to predict the transport properties of medicines such as human intestinal absorption, permeability to Caco-2 monolayer and permeation of blood-brain barrier [51]. Bioavailability results define the probability compounds to have minimum value of 10% oral bioavailability in rat or measurable Caco-2 permeability [52]. Considering the standard values, our four compounds pass through the Log S, bioavailability score, TPSA (Å^2^), Number of rotatable bonds value. The drug-likeness property experiment showed that our four compounds possess drug-like properties against tuberculosis.

Cytochrome P450 (CYPs) enzymes are metabolic enzymes, responsible for the biotransformation of ~90% FDA certified drugs [53]. The oxidative metabolism of drugs in phase I is achieved by the CYP system [54,55]. Nine of the isozymes under the CYP system scanned for the prediction of the metabolically vulnerable points using RS-WebPredictor tool [56]. It is a publicly available, web-based tool for predicting the regioselectivity of isozyme specific CYP mediated xenobiotic metabolism on any set of user-submitted molecules [57]. C22, C38, C21, C29 drugs displayed several sites of metabolism for CYP1A2, 2A6, 2B6, 2C8, 2C19, 2E1, 3A4, 2C9, 2D6 isoforms that suggested a satisfactory result.

In silico analysis of ADMET properties, absorption, distribution, metabolism, elimination and toxicity can help in determining the efficacy, pharmacological profile, mode of administration and safety of drugs. [58]. Human Intestinal Absorption (HIA) is a pharmacokinetic process that determines the effectivity of intestinal absorption or bioavailability of a drug upon oral administration, an anticipated route of drug administration [59]. Our four ligands showed a high rate of intestinal absorption and oral bioavailability. Caco-2 cell line, a prominent substitute to human intestinal epithelium (mucosa) is one of the in-vitro models to determine in vivo human intestinal absorption of drug molecules due to their morphological and functional resemblances with human enterocytes [60]. All of our probable drug candidates showed ability of penetrating Caco-2 cell line. P-glycoprotein (P-gp) is an efflux, ATP powered membrane-bound transport protein pump that is widely distributed throughout the body. It is a unique defensive barrier network that protects tissues from toxic xenobiotics by blocking entry to the cytosol and pumping its substrates to the exterior and also determines the uptake and distribution of pharmacotherapeutic drugs [61]. C22 acted as P-gp substrates as well as inhibitors. On the other hand, C38, C21, C29 were neither P-gp substrates nor P-gp inhibitors. Blood-Brain Barrier, a unique property possessed by the blood vessels in the Central Nervous System (CNS) tightly controls the transport of molecules, ions, nutrients and cells from blood to the brain and vice versa [62]. Consideration of BBB is an utmost priority for those drugs whose primary target is brain cells. Out of four ligands, only C38 was not capable of penetrating BBB.

As the incidence of anti-TB induced hepatotoxicity is one of the frequent causes behind the termination of anti-TB medication, assessing the potential hepatotoxicity of novel drugs is important [63]. As well as carcinogenicity test is necessary to classify a tumorigenic potentiality of drugs to assess the relevant risk in humans [64]. Only two ligands, C22 and C29 passed two of the criteria: - hepatotoxicity, and carcinogenicity as they showed negative result.

To validate conformational stability of our proposed drug upon binding with receptors, Molecular Dynamics Simulation was performed. Root Mean Square deviation (RMSD) analysis showed that binding of both control drugs and proposed drug C29 did not induce structural instability to the proteins because the reported RMSD change after ligand binding remained under 2 Å for both receptors. A thorough study of Root Mean Square Fluctuation (RMSF) curve revealed our tested compounds kept close contact with their active sites which was evident from their small range fluctuation under 1.5 Å and 2 Å for EthR and InhA complexes respectively. Though, because of loop regions on receptor proteins, greater fluctuation of RMSF was seen [65]. Our Radius of Gyration (Rg) analysis disclosed our proposed drug C29 and C22 induced fewer fluctuation of Rg upon binding with their respective receptors compared to the control drugs. This confirmed C29 and C22 did not cause instability to receptors. We compared our proposed drugs C29 and C22 with control drugs (Ethionamide and Isoniazid) through Molecular Dynamics Simulation (MDS), in which C29, C22 protein complexes represented stability throughout 10 ns simulation.

Starting from the preliminary docking analysis to Molecular Dynamics Simulation, our candidate compounds were thoroughly examined. At the stage of docking, our selected four candidates showed greater binding affinity in comparison to the control drugs. Additionally, the best four compounds examined for analyzing pharmacokinetics and pharmacodynamics properties and they showed promising results in drug-likeness and ADMET profiling. In all aspects, our chosen 3-[3-(4-Fluorophenyl)-1,2,4-oxadiazol-5-yl]-N-(2-methylphenyl)piperidine-1-carboxamide and 5-(4-Ethyl-phenyl)-2-(1H-tetrazol-5-ylmethyl)-2H-tetrazole showed satisfactory result than other two of the ligands. To conclude, 3-[3-(4-Fluorophenyl)-1,2,4-oxadiazol-5-yl]-N-(2-methylphenyl)piperidine-1-carboxamide and 5-(4-Ethyl-phenyl)-2-(1H-tetrazol-5-ylmethyl)-2H-tetrazole could be chosen as the most effective and promising candidates for as anti-TB drug.

## CONCLUSION

The number of people dying annually of TB is growing rapidly due to multiple drug resistance scenarios around the world. This situation demands newer anti-TB drugs to deal with the crisis. Drug repurposing is an easier and cheaper option to look for novel candidate as anti-TB drugs using different computational tools. The main objective of this study is to a find novel inhibitor against fast-mutating anti-TB drug targets. Our work incorporates pharmacore analysis, ADMET profiling, two-step molecular docking, followed by 10 ns Molecular Dynamics Simulation. Drugs with greater binding affinity than the control drugs are considered for determining Drug-likeness and ADMET analysis to evaluate their bioavailability and toxicity. Two of our screened compounds: 3-[3-(4-Fluorophenyl)-1,2,4-oxadiazol-5-yl]-N-(2-methylphenyl) piperidine-1-carboxamide and 5-(4-Ethyl-phenyl)-2-(1H-tetrazol-5-ylmethyl)-2H-tetrazole showed promising results with higher binding affinity with respective receptors and standard pharmacophoric properties. Molecular Dynamics Simulation study including RMSD, RMSF, Rg analysis confirmed their binding stability with respective proteins throughout the simulation timeline. Our present work can be productive finding new therapeutics against multiple drug resistant tuberculosis, having said that extensive in vitro and in vivo studies are required to prove our hypothetical analysis.

## ACKNOWLEDGEMENTS

The authors are thankful to Mohammad Mahfuz Ali Khan Shawan, the Assistant Professor of Dept. of Biochemistry and Molecular Biology, Jahangirnagar University, Savar, Dhaka for reviewing our research paper.

